# Physiological adaptation of sulfate reducing bacteria in syntrophic partnership with anaerobic methanotrophic archaea

**DOI:** 10.1101/2022.11.23.517749

**Authors:** Ranjani Murali, Hang Yu, Daan R. Speth, Fabai Wu, Kyle S. Metcalfe, Antoine Crémière, Rafael Laso-Pèrez, Rex R. Malmstrom, Danielle Goudeau, Tanja Woyke, Roland Hatzenpichler, Grayson L. Chadwick, Victoria J. Orphan

**Affiliations:** Division of Biology and Biological Engineering, California Institute of Technology, Pasadena, CA, USA; Division of Geological and Planetary Sciences, California Institute of Technology, Pasadena, CA, USA; Department of Physics and Astronomy, University of Southern California, Los Angeles, CA, USA; Max Planck Institute for Marine Microbiology, Bremen, Germany; ZJU-Hangzhou Global Scientific and Technological Innovation Center, Zhejiang, China; Department of Plant and Molecular Biology, University of California, Berkeley. Berkeley, CA, USA; National Center for Biotechnology, Madrid, Spain; DOE Joint Genome Institute, Department of Energy, Berkeley, CA, USA; Department of Chemistry and Biochemistry, Montana State University, Bozeman, MT, USA

## Abstract

Sulfate-coupled anaerobic oxidation of methane (AOM) is performed by multicellular consortia of anaerobic methanotrophic archaea (ANME) in obligate syntrophic partnership with sulfate-reducing bacteria (SRB). Diverse ANME and SRB clades co-associate but the physiological basis for their adaptation and diversification is not well understood. In this work, we explore the metabolic adaptation of four syntrophic SRB clades (HotSeep-1, Seep-SRB2, Seep-SRB1a and Seep-SRB1g) from a phylogenomics perspective, tracing the evolution of conserved proteins in the syntrophic SRB clades, and comparing the genomes of syntrophic SRB to their nearest evolutionary neighbors in the phylum Desulfobacterota. We note several examples of gain, loss or biochemical adaptation of proteins within pathways involved in extracellular electron transfer, electron transport chain, nutrient sharing, biofilm formation and cell adhesion. We demonstrate that the metabolic adaptations in each of these syntrophic clades are unique, suggesting that they have independently evolved, converging to a syntrophic partnership with ANME. Within the clades we also investigated the specialization of different syntrophic SRB species to partnerships with different ANME clades, using metagenomic sequences obtained from ANME and SRB partners in individual consortia after fluorescent-sorting of cell aggregates from anaerobic sediments. In one instance of metabolic adaptation to different partnerships, we show that Seep-SRB1a partners of ANME-2c appear to lack nutritional auxotrophies, while the related Seep-SRB1a partners of a different methanotrophic archaeal lineage, ANME-2a, are missing the cobalamin synthesis pathway, suggesting that the Seep-SRB1a partners of ANME-2a may have a nutritional dependence on its partner. Together, our paired genomic analysis of AOM consortia highlights the specific adaptation and diversification of syntrophic SRB clades linked to their associated ANME lineages.

## Introduction

Syntrophy is a form of metabolic cooperation between different microorganisms that enables the utilization of substrates which neither organism could metabolize on its own[1,2]. Microorganisms benefit from sharing nutrients and electrons in this way, combining their resources and allowing for less energy investment for each partner[1,3]. Syntrophic interactions appear to be specific in at least some cases, with the same organisms co-associating across different ecosystems and environments[4]. However, we do not yet understand the physiological basis driving the specificity of interactions, often because syntrophic associations are difficult to grow in the laboratory and characterizing the specificity of these interactions is challenging with uncultured syntrophic consortia in the environment[2]. A classic example of a syntrophic relationship exists between anaerobic methanotrophic archaea (ANME) and sulfate-reducing bacteria (SRB) in methane seeps[5–7]. ANME oxidize methane to CO_2_ anaerobically, in a geologically significant process known as anaerobic oxidation of methane(AOM) [8]. AOM is a thermodynamically unfavorable process that can only be completed when it is a coupled to an energetically favorable reaction such as sulfate reduction[9]. In multicellular consortia of ANME and SRB, coupling of methane oxidation to sulfate reduction appears to occur through direct interspecies electron transfer or DIET[10,11]. In addition to electron exchange, studies of syntrophic ANME-SRB consortia have also identified other hallmarks of syntrophy such as diazotrophic nitrogen exchange[12–14]. The ecophysiology of ANME/SRB consortia is complex and there are many divergent lineages of ANME and SRB that co-associate to form multicellular consortia. Given the diversity of interactions in ANME-SRB partnerships[14,15], there is a possibility that the basis and characteristics of some of these syntrophic partnerships may differ. Understanding the physiological basis of ANME-SRB interactions will provide insight into similar mechanisms that underpin the many syntrophic interactions in the biosphere.

Investigation of the archaeal and bacterial lineages involved in AOM identified at least three divergent taxonomic groups of archaea, by analysis of 16S rRNA gene sequences and fluorescence in situ hybridization (FISH) – ANME-1, ANME-2 and ANME-3[5,16,17]. All three of these groups are clades within the phylum Halobacterota. While ANME-1 is quite divergent and form a separate order called Methanophagales[18], all the ANME-2 clades, including ANME-2a (*Methanocomedenaceae*), ANME-2b (*Methanomarinus*), ANME-2c (*Methanogasteraceae*) and ANME-2d (*Methanoperedenaceae*) and members of the ANME-3 (*Methanovorans*), represent family or genus level taxa within the order Methanosarcinales[19]. Similar analyses of ANME-associated SRB revealed several clades of syntrophic SRB within the phylum Desulfobacterota: two clades related to Desulfobacterales (Seep-SRB1g and Seep-SRB1a)[14,20–22], seepDBB within the *Desulfobulbaceae*[23] and two more divergent groups - Seep-SRB2[24] and HotSeep-1[25]. ANME have also been reported to form spatial associations in consortia with additional microbial lineages including alpha- and beta-proteobacteria[26] and verrucomicrobia[27], Anaerolineales and Methanococcoides[28]. These partnerships and that of ANME with seepDBB, have however not been physiologically well-characterized and we thus do not include them in our analysis. Our analysis of ANME-SRB partnerships could be used to better under the nature of other associations between ANME and non-SRB species in the future. Previous research using FISH microscopy surveys[6,11,16,20,29,30] magneto-FISH[31,32], Bioorthogonal Non-canonical Amino Acid Tagging combined with fluorescence activated cell sorting (BONCAT-FACS)[27], and network analysis of statistical correlations in 16S rRNA gene amplicon sequencing data[14,33] revealed the most commonly observed partnerships between ANME and associated-SRB. Collectively, these results indicate that members of the ANME-1 order tend to partner with HotSeep-1 or Seep-SRB2 bacteria[24,25,33], ANME-2a with Seep-SRB1a[20,22], ANME-2b with Seep-SRB1g[14], ANME-2c with Seep-SRB1a[14,20] or Seep-SRB2[24,33], and ANME-3 with Seep-SRB1a[20] and some clades of *Desulfobulbaceae*[29]. These trends suggest that some lineages of ANME and syntrophic SRB partners, such as ANME-1, ANME-2c, Seep-SRB2 and Seep-SRB1a are capable of forming partnerships with multiple groups. However, greater taxonomic resolution within currently identified clades is required to test whether these partnerships truly are flexible (as is often true of syntrophies based on hydrogen exchange) or if subgroups can be identified that correspond to specific AOM syntrophic partnerships.

Metagenomics and comparative genomics have been used to identify metabolic traits unique to the previously identified ANME groups[15,19,34,35]. Consistent with their phylogenetic distance from ANME within the Methanosarcinales, members of the ANME-1 differ most from the others, as shown by distinctions in steps of the reverse methanogenesis pathway[34], their respiratory cytochromes *c*[19] and in their proposed use of quinone more often than methanophenazine as an electron carrier[19]. Significantly, the differences between the cytochrome *c* machinery of ANME-1 and that of ANME-2a/2b/2c and ANME-3 indicate that the mechanism of transferring electrons out of the cell must be very different even if it is ultimately still based on multi-heme cytochromes. Metabolic differences within ANME will directly affect the midpoint potential of the electron carrier that donates electrons to the syntrophic partner, their ability to fix and share nitrogen as well as their ability to synthesize and share other essential nutrients. As each of the syntrophic SRB are affected by the metabolic potential of their corresponding ANME partners and therefore, it is critical to consider the genomic traits of the SRB in the context of their syntrophic partners.

Among the syntrophic sulfate-reducing bacteria identified using 16S rRNA data and verified to be in partnership with ANME by FISH, genomes exist for members of the thermally adapted HotSeep-1 clade[24,25], as well as psychrophilic or mesophilic representatives of Seep-SRB1a[21,22], Seep-SRB1g[14,21] and Seep-SRB2[24]. These studies collectively confirmed that the complete pathway for dissimilatory sulfate reduction was universally present in all clades. Additionally, a large gene cluster containing multi-heme cytochromes (MHC) that is hypothesized to play a role in accepting electrons from ANME was detected in all genomes[21]. This previous work has established several defining characteristics of the syntrophic partners of ANME archaea. However, a robust taxonomic framework for identifying syntrophic SRB and differentiating them from their non-syntrophic SRB relatives is lacking, as is an evolutionary framework for understanding the metabolic adaptations in SRB that drove the formation of a syntrophic partnership with ANME. To bridge this knowledge gap, especially in light of the recent identification of a highly specific partnership between ANME-2b and Seep-SRB1g, we here employed comparative genomics to analyze the genomic traits of the dominant ANME associated SRB clades – Seep-SRB1a, Seep-SRB1g, Seep-SRB2 and HotSeep-1. Our analysis incorporated 576 bacterial genomes from the phylum Desulfobacterota from the GTDB database and 46 genomes of syntrophic SRB partners of ANME. This dataset included 15 previously unpublished metagenome-assembled genomes (MAGs) of syntrophic SRB and related clades recovered from seep sediments and mineral samples from three geographically distant locations in the Pacific Ocean including seeps located off the coast of Costa Rica, off the coast of S. California, as well as from hydrothermal vents in the Gulf of California. Importantly, several of these MAGs were sourced from fluorescent cell sorting and sequencing of single AOM consortia[27] enabling genomic analysis of these SRB syntrophs in the context of their specific ANME partner. Our analysis provides a framework for using genome-based phylogeny and 16S rRNA similarity to identify organisms from the four syntrophic SRB clades. In this work, we described a physiological framework comprising pathways that are important for the establishment of a syntrophic partnership between ANME and SRB and, with our phylogenomic framework, we identified multiple instances of metabolic adaptation that are specific to the syntrophic SRB, differentiating them from their nearest evolutionary neighbors. These instances were categorized as likely gene gains, losses, or specific cases of biochemical adaptation to a syntrophic lifestyle. With paired metagenomes of archaea and bacteria from single consortia, we also demonstrated that there appear to be specific instances of physiological adaptation of different Seep-SRB1a species to partnerships with different clades of ANME. Our study explored the diverse physiological strategies that underlie the different ANME-SRB partnerships, providing insight into the mechanism behind the establishment of the syntrophic partnership that is responsible for AOM.

## Results and Discussion

### Taxonomic diversity within syntrophic SRB of methanotrophic ANME

To investigate the adaptation of SRB to a partnership with ANME, we first placed them into their taxonomic context and assessed the phylogenetic diversity within the SRB clades (Seep-SRB1a, Seep-SRB1g, Seep-SRB2 and HotSeep-1). For this analysis we compiled a curated dataset of metagenome assembled genomes (MAGs) from these SRB clades including 34 previously published genomes[21,22,25,36–40] and 12 MAGs assembled for this study. Five of these genomes were reconstructed from seep samples collected off the coast of California, Costa Rica, and within the Gulf of California. We also sequenced single ANME-SRB consortia that were sorted by FACS (Fluorescence-activated cell sorting) after they were SYBR-stained as previously described[28]. With this technique, we could be confident of the assignment of partners that physically co-associate within the sequenced aggregates and begin to identify partnership specific characteristics. From sequencing of single consortia, we obtained 2 genomes of ANME-2b associated Seep-SRB1g, 1 genome of ANME-2a associated Seep-SRB1a and 3 genomes of ANME-2c associated Seep-SRB1a **(Table 1).** We recovered an additional 3 genomes of the nearest evolutionary neighbors of HotSeep-1 within the order Desulfofervidales since this order of bacteria is very poorly represented in public databases. Our dataset for comparative genomics analysis comprised the above mentioned 46 genomes of syntrophic SRB and 576 other bacteria from Desulfobacterota. Having compiled this dataset of syntrophic SRB, we also designated type material and proposed formal names for three of the syntrophic SRB clades, Seep-SRB2 (*Candidatus* Desulfomithrium gen. nov.), Seep-SRB1a (*Candidatus* Syntrophophila gen. nov.), Seep-SRB1g (*Candidatus* Desulfomellonium gen. nov.). The genomes designated as type material are identified in **Figure 1** and **Supplementary Figure 1**. Further details are available in **Supplementary information**.

**Figure 1.**
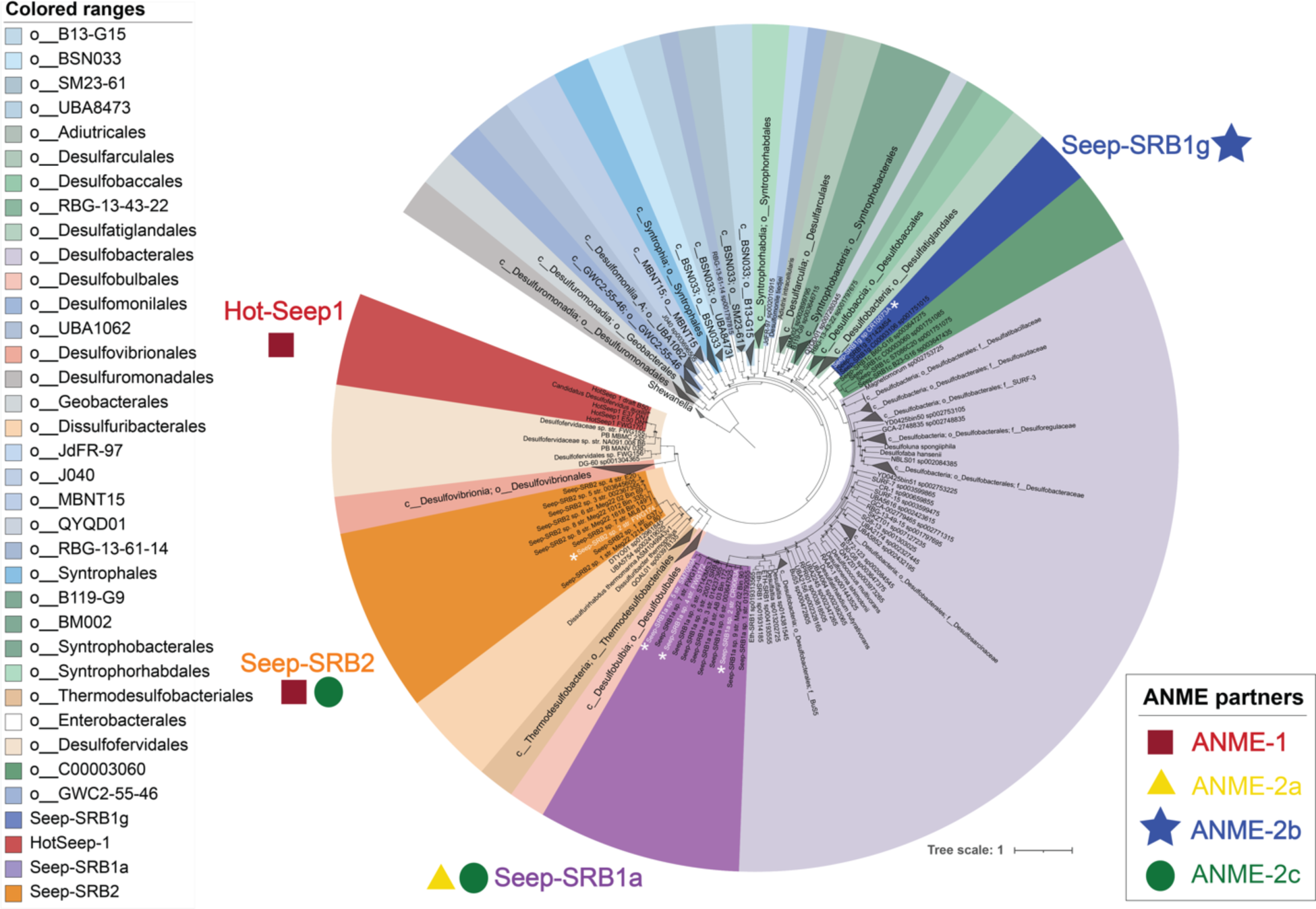
Taxonomic diversity of syntropic sulfate reducing bacteria. A concatenated gene tree of 71 ribosomal proteins from all the Desulfobacterota genomes within the GTDB database release 95 was made using Anvi’o[1]. Genomes from the genus *Shewanella* were used as outgroup. Within this tree, the four most common lineages of the syntrophic partners of anaerobic methanotrophic archaea (ANME) – Seep-SRB1a, Seep-SRB1g, Seep-SRB2 and Hot-Seep1 are highlighted. While Seep-SRB1a is a genus within the order Desulfobacterales, Seep-SRB1g and Seep-SRB1c together appear to form a closely related order-level taxonomic clade within the class Desulfobacteria. Seep-SRB2 is a genus within the order Dissulfuribacterales while Hot-Seep1 is its own species within the order Desulfofervidales. The proposed type strains are identified on the tree in white with a white asterisk adjacent to the label.

Details for the phylogenetic placement of each of these clades using 16S rRNA phylogeny, concatenated ribosomal protein phylogeny and the Genome Taxonomy Database are provided in **Materials and Methods** and **Supplementary Information**. HotSeep-1 is a species within the order Desulfofervidales, an order that is largely associated with thermophilic environments (with one exception, Desulfofervidales sp. DG-60 was sequenced from the White Oak Estuary[41]). Members of HotSeep-1 are the best characterized members of this order and are known to be syntrophic partners to thermophilic clades of methane-oxidizing ANME-1[15,24] as well as alkane-oxidizing archaeal relatives ‘*Candidatus* Syntrophoarchaeum butanivorans’, ‘*Candidatus* Syntrophoarchaeum caldarius’[42] and ethane-oxidizing ‘*Candidatus* Ethanoperedens thermophilum’[36]. Seep-SRB2 is a genus level clade within the order Dissulfuribacterales[43–45] and class Dissulfuribacteria. Dissulfuribacterales include the genera Dissulfuribacter and Dissulfurirhabdus[44,45], which are chemolithoautotrophs associated with sulfur disproportionation. Seep-SRB1g is a species level clade which groups within a taxonomic order that also includes Seep-SRB1c (**Figure 1, Table 1**). This order falls within the class Desulfobacteria along with the sister order Desulfobacterales. Like the Desulfofervidales, the order with Seep-SRB1g is poorly characterized, yet its most well-described members are the Seep-SRB1g that are obligate syntrophic partners of ANME, accepting electrons from the archaeal partner to reduce sulfate[14,21]. Seep-SRB1a is a genus level clade that along with the genus Eth-SRB1 forms a distinct family within the order Desulfobacterales (**Figure 1**, **Supplementary Figure 1**, **Supplementary Table 2**). Many of the well-characterized members of Desulfobacterales such as *Desulfococcus oleovorans*, *Desulfobacter hydrogenophilus*, *Desulfosarcina* BuS5 are known as hydrogenotrophs and hydrocarbon degraders[46–48]. The nearest evolutionary relative of Seep-SRB1a are the Eth-SRB1 first characterized as a syntrophic partner of ethane-degrading archaea[49]. Each of the four syntrophic SRB clades have evolved from taxonomically divergent ancestors with different metabolic capabilities. While the adaptation to a syntrophic partnership with ANME appears to have been convergently evolved in these clades, their evolutionary trajectories are likely to be different.

Species diversity within each of these clades was inferred by calculating the average nucleotide identity (ANI) (**Supplementary Figure 1**) and 16S rRNA sequence similarity (**Supplementary Table 2**) between different organisms that belong to each clade, using a 95 % ANI value and 98.65 % similarity in 16S rRNA as cut-offs to delineate different species. Partnership associations, as identified in previous research by our group and others, by FISH[20,24,25], magneto-FISH[32] or FACS sorting[27] and single-aggregate sequencing[28] are depicted in **Figure 1** and **Supplementary Figure 1** with further details provided in **Supplementary Information**. All the genomes of Seep-SRB1g in our curated database belong to one species-level clade and thus far, have been shown to partner only ANME-2b[14]. In contrast, there is greater species diversity within the clades that are known to partner more than one clade of ANME, Seep-SRB2 and Seep-SRB1a. Whether this diversification is driven by adaptation to partnerships with multiple ANME clades remains to be seen. This pattern is also not consistent with HotSeep-1, a species level clade that partners multiple archaeal species. A better understanding of the physiological basis for syntrophic partnership formation in each of these clades will provide a framework to understand their unique diversification trajectories.

### Comparative genome analysis of syntrophic SRB

To develop insight into the adaptation of SRB to syntrophic partnerships with ANME, we used a comparative genomics analysis approach to 1) identify the unique features of known syntrophic SRB partners relative to their closest non-syntrophic relatives and 2) compare the physiological traits that define the diversity within a given taxonomic clade of syntrophic partner bacteria. For our first objective, we placed the metabolic traits of SRB into the phylogenetic context of the Desulfobacterota phylum, correlating the presence or absence of a physiological trait within the context of genus, family and order level context of each syntrophic SRB clade. As an example, we demonstrate that the multi-heme cytochrome conduit[21] implicated in DIET between ANME and SRB is rare in non-syntrophic Desulfobacterota suggesting that this trait is part of a required adaptation for this syntrophic relationship (**Figure 2**).

**Figure 2.**
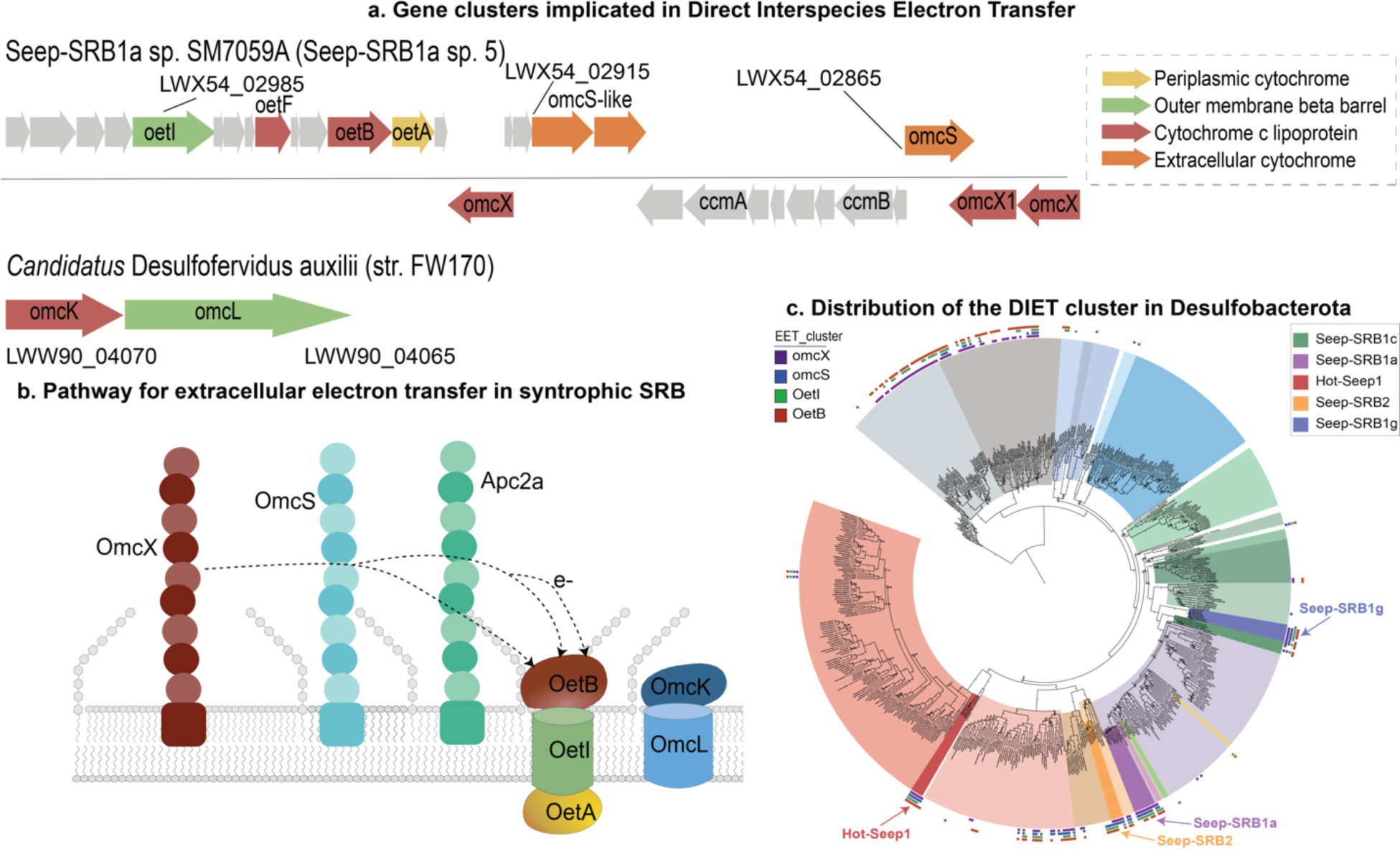
Gene organization and distribution of the putative cluster implicated in direct interspecies electron transfer from syntrophic sulfate reducing bacteria. a. The syntenic blocks of genes implicated in direct interspecies electron transfer including the putative extracellular electron transfer conduit OetABI, and the operon encoding for OmcKL from HotSeep-1 (*Candidatus* Desulfofervidus auxilii). b. A model of the putative extracellular electron transfer within syntrophic SRB. ANME electrons are likely to be accepted by one of three putative nanowires formed by multi-heme cytochromes homologous to OmcX, OmcS and a cytochrome we named Apc2a. The electrons from this nanowire would then be transferred to the porin:multiheme cytochrome *c* conduits formed by OetABI or OmcKL, and ultimately to different periplasmic cytochromes *c*. c. The distribution of the putative DIET cluster in the phylum Desulfobacterota is mapped onto a whole genome phylogenetic tree of Desulfobacterota based on the presence of OmcX, OetI, OetB, and Apc2a. This cluster is not widely found except in the orders Desulfuromonadales and Geobacterales, and the classes Desulfobulbia and Thermodesulfobacteria.

We also investigated the physiological differences between the species of each syntrophic SRB clade. Two of the syntrophic SRB clades, Seep-SRB1g and HotSeep-1 have low diversity, with representatives from different seep and vent ecosystems each belonging to a single species-level clade. The clades Seep-SRB2 and Seep-SRB1a in contrast, contain multiple species. To better understand the genomic features underlying this diversity, we performed a comparative analysis of species within the Seep-SRB1a and Seep-SRB2 to identify conserved genes across the clade and species-specific genes. A detailed description of the analysis methods is available in **Materials and Methods** and **Supplementary Information** (**Supplementary Figures 5, 6, Supplementary Tables 3, 4)**. For this comparative analysis, we primarily focused on pathways that are predicted to be important for the syntrophic interactions between ANME and SRB. In the following section, we describe the pathways within the syntrophic SRB in greater detail and their significance for a syntrophic lifestyle – extracellular electron transfer, membrane-bound electron transport chain, electron bifurcation, carbon fixation, nutrient sharing, biofilm formation, cell adhesion and partner identification. Lastly, we explicitly compare the losses and gains of the genes encoding for the above pathways across the syntrophic SRB and infer the evolutionary trajectory of adaptation towards a syntrophic partnership.

### Respiratory pathways in the four syntrophic SRB clades demonstrate significant metabolic flexibility

The respiratory pathways in syntrophic SRB are defined by the necessity of ANME to transfer the electrons derived from methane oxidation to SRB. These electrons are then transferred across the outer membrane to periplasmic electron carriers. These periplasmic electron carriers donate electrons to inner membrane complexes and ultimately, to the core sulfate reduction pathway. Some of the electrons are also used for assimilatory pathways such as carbon fixation. Accordingly, our analysis of the respiratory pathways is split into a description of the pathways for interspecies electron transfer, electron transfer across the inner membrane, and carbon fixation pathways. In several instances we also note the potential for multiple complexes having redundant functionality which may afford respiratory flexibility within these pathways and emphasize the steps or reactions at which energy conservation likely occurs.

#### a. Multiple pathways for interspecies electron transfer between ANME and SRB

The dominant mechanism of interspecies electron transfer between ANME and SRB was proposed to be direct interspecies electron transfer (DIET). This hypothesis is supported by the presence of multi-heme cytochromes in genomes of ANME-2a, 2b and 2c[11], the presence of nano-wire like structures that extend between ANME-1 and its partners Hot-Seep1[10] and Seep-SRB2[24], and the presence of hemes in the extracellular space between archaeal and bacterial cells in ANME-SRB aggregates[11,24]. This hypothesis was also supported by the presence of a putative large multi-heme cytochrome:porin type conduit, analogous to the conduits in *Geobacter sp.*[50] and other gram-negative bacteria that have been shown to participate in extracellular electron transfer (EET)[50], in Seep-SRB1g[21], Seep-SRB2[24] and Hot-Seep-1[10]. Our analysis of a more comprehensive dataset of syntrophic bacterial genomes confirms the presence of this porin:cytochrome *c* conduit in all the four syntrophic bacterial clades studied (**Supplementary Table 5**). Henceforth, we refer to this as the as the (**O**uter-membrane bound **e**xtracellular electron **t**ransfer) or Oet-type conduit (OetI was verified to be capable of EET, data not shown). This conduit includes a periplasmic cytochrome *c* (OetA), an outer-membrane porin (OetI), and extracellular facing cytochrome *c* lipoprotein (OetB) (**Figure 2b, Figure 3**). The OetI-type conduit was first identified in *G. sulfurreducens* and is expressed when a Geobacter mutant of omcB is grown on Fe(III) oxide[51]. The oetABI cassette is found in all four syntrophic SRB clades, and often includes two or three other putative extracellular cytochromes *c*, including homologs of OmcX[21], OmcS (Supplementary alignment MSA1) and a 6-heme cytochrome that we termed apc2a (**Supplementary Table 5**). If they are not found as part of the oet cluster, they could be found elsewhere on the genome, possibly due to genomic rearrangement after acquisition of the cassette (**Supplementary Tables 6, 7**). The omcX and omcS-like genes in the oet gene cassette are often found in an analogous position to omcS and omcT in G. *sulfurreducens* (**Figure 2**). Based on the homology of one of the cytochromes to OmcS, which polymerizes to form long and highly conductive filaments that facilitate extracellular electron transfer in *Geobacter*[52], we propose that the extracellular cytochromes *c* in this gene cassette perform a similar function, forming filaments that accept electrons from ANME. This is consistent with heme staining of the intercellular space between ANME and SRB, and the observation of filaments that connect the partners[11,24]. This is also consistent with the fact that different extracellular cytochromes are amongst the most highly-expressed proteins in the syntrophic SRB: ANME-1/Seep-SRB2[24] (OmcX, OmcS-like and apc2a), ANME-1/HotSeep-1[24] (OmcX and OmcS-like), ANME-2c/Seep-SRB2[24] (OmcX) aggregates and ANME-2a/Seep-SRB1a[22] (OmcX, OmcS-like). The presence of multiple copies of these putative filament forming proteins in the syntrophic SRB genomes is indicative of their importance to the physiology of syntrophic SRB. The mechanism of electron transfer from extracellular cytochrome filaments to the interior of the cells in *Geobacter* is not well understood. However, a porin:cytochrome *c* conduit is always expressed under the same conditions as a cytochrome *c* containing filament in *Geobacter* (omcS along with extEFG or omcABC under Fe (III) oxide reducing conditions and omcZ along with extABCD during growth on an electrode[53]) and in ANME-SRB consortia (OmcS/OmcX with OetABI or OmcKL). These findings suggest that each cytochrome *c* filament could act in concert with a porin:cytochrome *c* conduit (**Figure 2**) to transfer electrons from the extracellular space to the periplasm.

**Figure 3.**
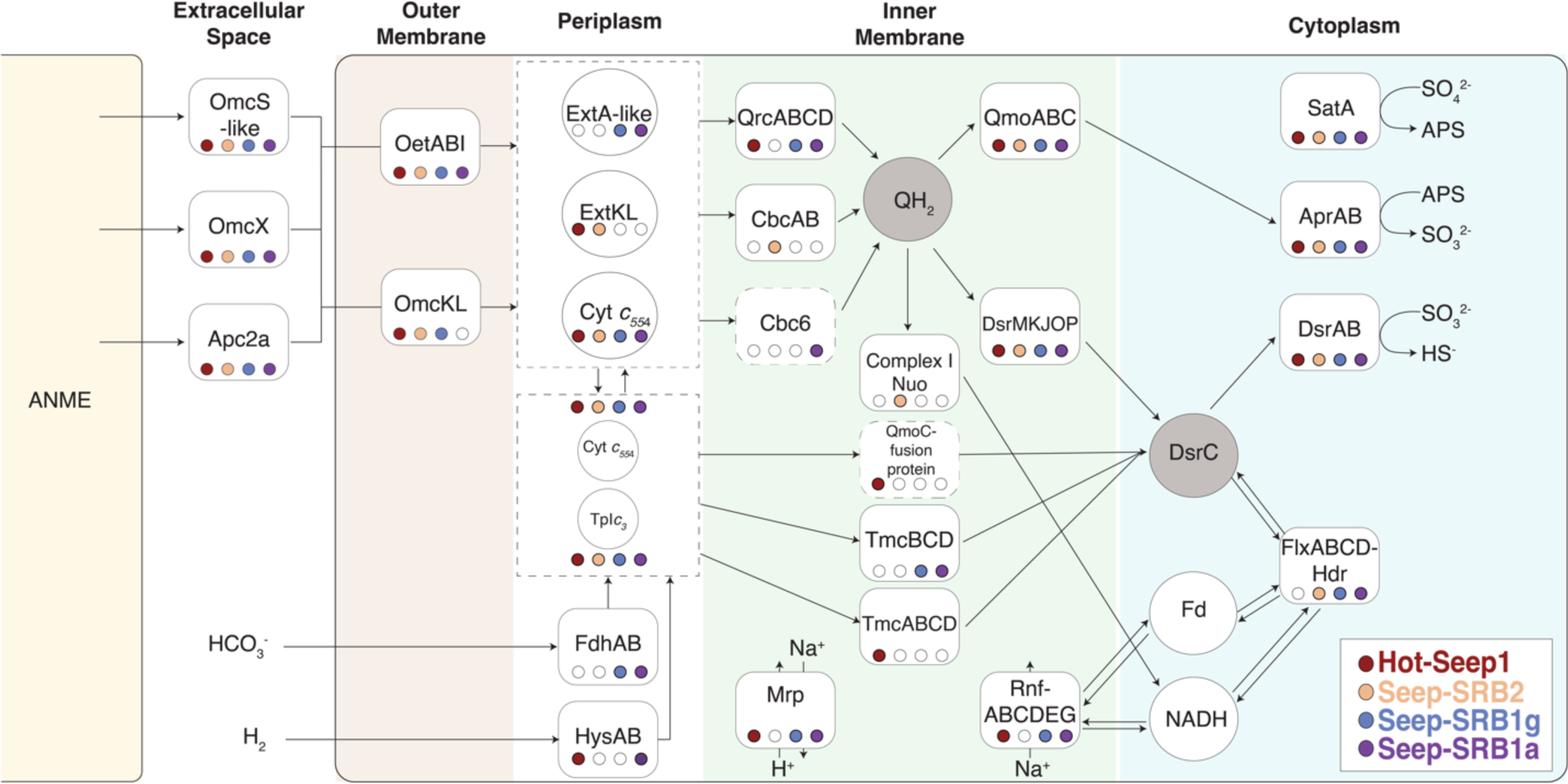
Summary of the different electron transport chains in syntrophic sulfate reducing bacteria. The various respiratory proteins essential for the electron transport chain within the syntrophic SRB are identified and marked within their predicted cellular compartments. Filled circles indicate their presence in each of the four syntrophic sulfate reducing bacterial clades, HotSeep-1 (red), Seep-SRB2 (orange), Seep-SRB1g (blue) and Seep-SRB1a (purple). The two typical acceptors of electrons transferred across the inner membrane, quinols (QH_2_) and DsrC, are indicated in shaded circles. These are the two nodes which much of the respiratory flexibility of the syntrophic SRB revolves around.

While oetABI is conserved in all four syntrophic SRB clades, there are two other putative porin:cytochrome *c* conduits in syntrophic SRB. A porin (HS1_RS02765) and extracellular cytochrome c (HS1_RS02760) homologous to OmcL and OmcK from *G. sulfurreducens* is found in HotSeep-1 (**Supplementary Table 6, 7**) and expressed at a four-fold higher level than the oetABI conduit[24]. OmcK and OmcL were also upregulated in *G. sulfurreducens* when it is grown on hematite and magnetite[54]. There is no gene encoding a periplasmic cytochrome *c* adjacent to these genes and this is unusual for previously characterized EET conduits but, given the large number of periplasmic cytochromes in HotSeep-1, it is conceivable that another cytochrome *c* interacts with the OmcL/K homologs. This conduit is also found in Seep-SRB2 sp. 1, 2, 7 and 8 but does not appear to be expressed as highly as the OetABI in the ANME-1/Seep-SRB2 consortia[24]. A different putative conduit including the porin, extracellular and periplasmic cytochromes *c* is present in the Seep-SRB1g genomes (LWX52_07950-LWX52_07960)(**Supplementary Table 6, 7**). This conduit does not have identifiable homologs in *Geobacter.* The presence of multiple porin:cytochrome *c* conduits in the syntrophic partners suggests some flexibility in use of electron donors, possibly from different syntrophic partners. For HotSeep-1, this observation is consistent with its ability to form partnerships with both methane and other alkane-oxidizing archaea[55]. The role of the second conduit is less clear in Seep-SRB1g which to date has only been shown to partner with ANME-2b, and currently lacks representation in enrichment cultures[14]. Future investigation of the multiple syntrophic SRB extracellular electron transfer pathways and the potential respiratory flexibility it affords to their partner archaea using transcriptomics, proteomics and possibly heterologous expression methods will further expand our understanding of electron transfer in these diverse consortia.

While direct interspecies electron transfer is believed to be the dominant mechanism of syntrophic coupling between the ANME and SRB partners, the potential to use diffusible intermediates such as formate and hydrogen exists in some genomes of syntrophic SRB. Hydrogenases are present in HotSeep-1, which can grow without ANME using hydrogen as an electron donor[25]. We also identified periplasmic hydrogenases in Seep-SRB1a sp. 1, 5 and 8 (**Supplementary Tables 7**) which suggest that these organisms could use hydrogen as an electron donor. However, in Seep-SRB1a these hydrogenases are expressed at low levels (less than a twentieth of the levels of DsrB) in the ANME-2a/Seep-SRB1a consortia[22]. Further, previous experiments showed that the addition of hydrogen to ANME-2/SRB consortia did not inhibit anaerobic oxidation of methane suggesting that hydrogen is not the predominant agent of electron transfer between ANME and SRB[26,56]. Perhaps, hydrogenases are used by Seep-SRB1a to scavenge small amounts of hydrogen from the environment. While membrane bound and periplasmic hydrogenases are present in non-syntrophic Seep-SRB1c (**Supplementary Table 7**), no hydrogenases are found in the syntrophic relative of Seep-SRB1c and ANME partner, Seep-SRB1g. Similarly periplasmic hydrogenases are present in Dissulfuribacteriales and absent in Seep-SRB2 (one exception in 18 genomes), suggesting that in both these partners, the loss of periplasmic hydrogenases is part of the adaptation to their syntrophic partnership with ANME. We also identified periplasmic formate dehydrogenases in Seep-SRB1g and Seep-SRB1a sp. 2, 3, 8, 9 (**Supplementary Table 7**). The periplasmic formate dehydrogenase from Seep-SRB1g is expressed in the environmental proteome at Santa Monica Mounds[21], but no transcripts from the formate dehydrogenases of Seep-SRB1a were recovered in the ANME-2a/Seep-SRB1a incubations[22]. It is possible that these syntrophic SRB scavenge formate from the environment. Alternatively, a recent paper found a hybrid of electron transfer by DIET and by diffusible intermediates (mediated interspecies electron transfer or MIET) to be energetically favorable[57]. In this model, the bulk of electrons would still be transferred by DIET, but up to 10 % of electrons could be shared by MIET via formate[57], an intermediate suggested in earlier studies[26,56]. This might be possible in ANME/SRB consortia with HotSeep-1, some species of Seep-SRB1a and Seep-SRB1g, but not in consortia with Seep-SRB2. The absence of periplasmic formate dehydrogenases and hydrogenases in Seep-SRB2 as previously observed[24] is also true in our expanded dataset. If a diffusive intermediate should play a role in mediating electron transfer between ANME-2c or ANME-1 and Seep-SRB2, it is not likely to be formate or hydrogen.

#### b. Different pathways for electron transfer across the inner membrane in syntrophic SRB

Multiheme cytochromes *c* in SRB are known to mediate diverse modes of electron transfer from different electron donors to a conserved sulfate reduction pathway[58]. There is significant variety in the number and types of cytochromes *c* present in sulfate reducing bacteria from the phylum Desulfobacterota[58] and an even greater number of large cytochromes is present in syntrophic SRB[21,24]. To explore the potential for different routes of electron transfer, we performed an analysis of all cytochromes *c* containing four hemes or more from the genomes of syntrophic SRB (see Materials and Methods) and identified at least 27 different types of cytochromes *c.* We split these cytochromes *c* into those predicted to be involved in extracellular electron transfer, those that act as periplasmic electron carriers and those that are components of protein complexes involved in electron transfer across the inner membrane (**Supplementary Table 6, Supplementary Information**). Conserved across the syntrophic SRB partners of ANME were the cytochromes forming the core components of the EET pathway– OetA, OetB, OmcX and OmcS-like and Apc2a extracellular cytochromes, and two periplasmic cytochromes of the types, TpI*c_3_*[59] and cytochrome *c_554_*[58,60]. Beyond the conserved periplasmic cytochromes *c,* TpI*c_3_* and cytochrome *c_554,_* there are also cytochromes binding 7-8 hemes that are unique to different SRB clades (**Supplementary Table 6**). These include a homolog of ExtKL[61] from *G. sulfurreducens* that is highly expressed in Seep-SRB2 spp. 1 and 4 during growth in a syntrophic partnership with ANME[24], and a homolog of ExtA from *G. sulfurreducens*[50] protein expressed in the ANME-2a/Seep-SRB1a consortia[22]. Previous research has suggested that the tetraheme cytochromes *c* are not selective as electron carriers and play a role in transferring electrons to multiple different protein complexes[62]. It is possible that these larger 7-8 heme binding cytochromes *c* have a more specific binding partner. Both the ExtKL and ExtA-like proteins are very similar (over 45% sequence similarity) to their homologs in Desulfuromonadales. Since the OetI-type conduit is also likely transferred from this order, they might act as binding partners.

In SRB, the electrons from periplasmic electron donors (reduced by DIET or MIET) are delivered through inner membrane bound complexes to quinones or directly to the heterodisulfide DsrC in the cytoplasm via transmembrane electron transfer[58] (**Figure 3**). The electrons from quinones or DsrC are ultimately used for the sulfate reduction pathway (including SatA, AprAB and DsrAB)[58,21,24]. Two conserved protein complexes are always found along with this pathway – the Qmo complexes transfers electrons from reduced quinones to AprAB and the DsrMKJOP complexes transfers electrons from quinones to DsrC and through DsrC to DsrAB. Since both these complexes use electrons from reduced quinones, the source of reduced quinones in the inner membrane is critical to different sulfate respiration pathways. The quinol reducing complexes and complexes that reduce DsrC provide respiratory flexibility to sulfate reducing bacteria. We also note here that the reduction of AprAB coupled to the oxidation of menaquinone is expected to be endergonic. There is a proposal that QmoABC might function through flavin-based electron confurcation (FBEC), using electrons from reduced quinones and a second electron donor such as ferredoxin to reduced AprAB[63], Since, it is not clear what the electron donor is likely to be, we do not explicitly consider this reaction in our analysis. A summary of all the putative complexes that are involved in the electron transport chains of the four syntrophic SRB is visualized in **Figure 3** to detail how electron transport pathways vary among the clades. A more detailed list of complexes present is found in **Supplementary Tables 6-8**. The respiratory pathways in HotSeep-1, Seep-SRB1a and Seep-SRB1g are broadly similar in structure and are predicted to use the Qrc complex to transfer periplasmic electrons to the quinone pool and Tmc to reduce cytoplasmic DsrC. Their pathways are analogous to the respiratory pathways in *Desulfovibrio alaskensis*[59,64]. Qrc is the major site of energy conservation in this respiratory pathway. Protons are translocated by Qrc from the cytoplasmic side to the periplasmic active side. This movement of charges across the membrane leads to the generation of proton motive force (pmf) that can be utilized by ATP synthase to generate ATP[65]. In Seep-SRB2, Qrc is absent and we hypothesize that CbcBA a protein complex that appears to be horizontally transferred from the Desulfuromonadales (**Figure 3, Figure 4**) mediates electron transfer between periplasmic cytochromes *c* and quinones[66]. This is supported by the fact that CbcBA is highly expressed during AOM between ANME-1/Seep-SRB2 and ANME-2c/Seep-SRB2[24]. In *Geobacter sulfurreducens*, which also does not have Qrc this cytoplasmic membrane-bound oxidoreductase is expressed during growth on Fe(III) at low potential and is important for iron reduction and growth on electrodes at redox potentials less than -0.21 mV[66]. During AOM, the CbcBA protein in Seep-SRB2 is predicted to run in the reverse direction, reducing quinols using electrons from DIET electrons supplied by ANME archaea as opposed to functioning in the metal reducing direction. While the reversibility of this complex has not been biochemically established, the high levels of expression of this complex suggest that this is likely functional in the electron transport chain of Seep-SRB2. It is not clear what the likely site of energetic coupling is within the Seep-SRB2 respiratory chain. In the absence of the Qrc complex, the most likely mechanism for energetic coupling might exist through the action of a Q-loop mechanism[67]. In this mechanism, energy is conserved by the combined action of two protein complexes that reduce and oxidize quinols, leading to the uptake and release of protons on opposite sides of the cytoplasmic membrane. The Q-loop mechanism in Seep-SRB2 would likely involve CbcBA and a quinol oxidizing complex such as Qmo.

**Figure 4.**
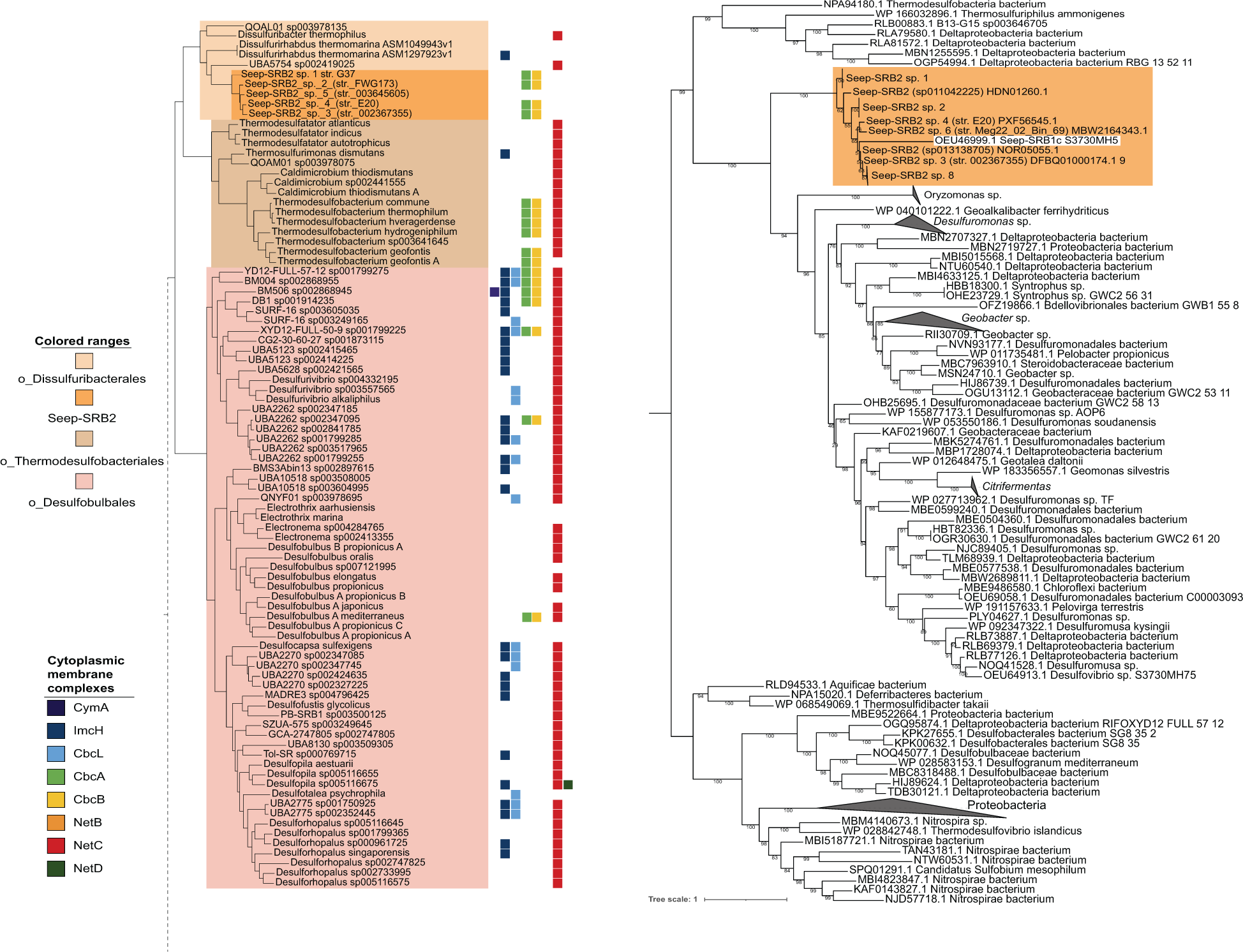
*cbcBA* as an example of horizontal gene transfer events into Seep-SRB2. We demonstrate one example of an important gene transfer event involving a metabolic gene. The function of cbcBA is essential for the central respiratory pathway in Seep-SRB2 and this gene was acquired by horizontal gene transfer. A. The presence of CymA, CbcL, CbcAB and NetBCD, commonly used electron donors to the extracellular electron transfer (EET) conduits in *Shewanella* and *Geobacter* are mapped on to the classes Thermodesulfobacteria and Desulfobulbia. B. CbcB protein sequences were aligned using MUSCLE[2] and then a phylogenetic tree was inferred using IQ-Tree2[3]. The CbcB sequences from Seep-SRB2 are highlighted in orange.

In addition to the most likely pathways of electron transfer in the syntrophic SRB, as established using transcriptomic data on ANME/SRB partnerships[22,24] (**Figure 2**), other inner membrane complexes exist in these genomes that may provide additional respiratory flexibility. HotSeep-1 genomes contain a complex that involves an HdrA subunit and a QmoC-fusion protein that also binds hemes *c*. This complex would likely transfer electrons from cytochromes *c* to the DsrC (AMM42179.1-AMM42180.1). The presence of HdrA might indicate a role in electron bifurcation by this complex. It is highly expressed during methane oxidation conditions in the ANME-1/HotSeep-1 consortia to a fifth of the level of the Tmc complex that would play a similar role in electron transfer[24]. In some Seep-SRB1a and Seep-SRB1g genomes, there is a homolog of Cbc6 (LWX51_14670-LWX51_14685) identified in *Geobacter*[68] and implicated in electron transfer from periplasmic cytochromes *c* to the quinol pool. A NapC/NirT homolog[69] was conserved in Seep-SRB1g (OEU53943.1-OEU53944.1) and some Seep-SRB2, and another conserved complex that includes a cytochrome *c* and ruberythrin (AMM39991.1-AMM39993.1) is present in Seep-SRB1g and HotSeep-1. Further research is needed to test whether there are conditions under which these complexes are expressed. Our analysis indicates some degree of respiratory specialization in the syntrophic SRB genomes such as the loss of hydrogenases in Seep-SRB1g and Seep-SRB2 compared to their nearest evolutionary neighbors, suggesting an adaptation towards a partnership with ANME. However, considerable respiratory flexibility still exists within the genomes of these syntrophic partners as is suggested by the presence of the formate dehydrogenases in Seep-SRB1g and Seep-SRB1a, multiple EET conduits in HotSeep-1 and Seep-SRB2 and multiple inner membrane complexes in Seep-SRB1a and HotSeep-1.

#### c. Cytoplasmic redox reactions, electron bifurcation and carbon fixation

The electron transport chain outlined above would transfer electrons from periplasmic cytochromes *c* to the cytoplasmic electron carrier DsrC or directly to the sulfate reduction pathway. However, the electron donors for carbon or nitrogen fixation are typically NADH, NADPH or ferredoxin[70]. The transfer of electrons from DsrC to these reductants likely happens through the action of membrane-bound Rnf and Mrp[71–73] in Seep-SRB1a, Seep-SRB1g and Hot-Seep1. In marine environments, the naturally occurring sodium gradient can be used to generate ferredoxin from NADH or vice versa using the Rnf complex, while the Na^+^/H^+^ antiporter, Mrp, can transport Na^+^ or H^+^ in response to the action of Rnf[74]. The ferredoxin generated from this process can then be used for assimilatory pathways. In Seep-SRB2, which does not contain Rnf or Mrp (**Supplementary Figure 9**), the NADH needed for carbon fixation is likely obtained through the oxidation of quinol by complex I and the dissipation of proton motive force. In addition to Complex I or Rnf and Mrp, there are additional cytoplasmic protein complexes that can recycle reducing equivalents between DsrC, ferredoxin and NADH. One of these protein complexes is electron bifurcating Flx-Hdr[75,76] which can oxidize two molecules of NADH to reduce one molecule of ferredoxin and one molecule of DsrC. Several putative oxidoreductase complexes in the syntrophic SRB genomes are compiled in **Supplementary Table 7** and **Supplementary Figures 10, 11**.

Syntrophic sulfate-reducing members of the Seep-SRB1a, Seep-SRB1g, and Seep-SRB2 have been shown to fix carbon using the Wood-Ljungdahl pathway, while organisms of the clade HotSeep-1 partnering with ANME-1 are predicted to fix carbon using the reductive tricarboxylic acid cycle (rTCA)[21,24,70]. Analysis of gene synteny for a number of Seep-SRB1a, Seep-SRB1g and Seep-SRB2 MAGs uncovered a number of heterodisulfide (HdrA) subunits and HdrABC adjacent to enzymes involved in the Wood-Ljungdahl pathway (**Figure 10**). These subunits are typically implicated in flavin-based electron bifurcating reactions utilizing ferredoxins or heterodisulfides and NADH[72], The prevalence of such HdrA containing complexes is also common in ANME[15]. Specifically, Seep-SRB1g has an HdrABC adjacent to metF that is predicted to encode for a putative metF-HdrABC, performing the reduction of methylene tetrahydrofolate reductase coupled to the endergonic reduction of ferredoxin to NADH, the same reaction the bifurcating metFV-HdrABC described below. In Seep-SRB1g, there are also two copies of HdrABC next to each other whose function requires further analysis (**Supplementary Figure 10**). These complexes are absent in the related group Seep-SRB1c, a lineage which has not yet been found in physical association with ANME (**Supplementary Figure 10**). The presence of electron bifurcation machinery in the carbon fixation pathways within several syntrophic SRB lineages, suggests that they are optimized to conserve energy (**Supplementary Figure 10**). This is reminiscent of the MetFV-HdrABC in the acetogen *Moorella thermoacetica*[72] in which the NADH-dependent methylene tetrahydrofolate reductase reaction within the central metabolic pathway is coupled to the endergonic reduction of ferredoxin by NADH, allowing for the recycling of reducing equivalents. Members of the Seep-SRB1g also have a formate dehydrogenase (fdhF2) subunit adjacent to nfnB, the bifurcating subunit of nfnAB, which performs the NADPH-dependent reduction of ferredoxin (**Supplementary Figure 11**). This complex is predicted to function as an additional bifurcating enzyme that would allow for the recycling of NADPH electrons. In addition, HotSeep-1, Seep-SRB2 and Seep-SRB1g appear to have homologs of electron transfer flavoproteins, etfAB, that are expected to be electron bifurcating. These homologs of etfAB cluster with the previously identified bifurcating etfAB, and possess the same sequence motif that was previously shown to correlate with the electron bifurcating etfAB[77] (**Supplementary Table 7**). While the capability of electron bifurcation by these enzyme complexes needs to be biochemically confirmed, the possibility of a high number of bifurcating complexes, especially those connected to the carbon fixation pathway, in the genomes of syntrophic SRB partners of ANME is compelling. It could be argued that this is a natural adaptation to growth in very low energy environments or to low-energy metabolism. In fact, some of these complexes are present in other bacteria of the order Desulfofervidales and genus Eth-SRB1. These adaptations could provide an additional energetic benefit for the syntrophic lifestyle, itself an adaptation to low-energy environments.

#### d. Cobalamin auxotrophy and nutrient sharing in syntrophic SRB

Research on the AOM symbiosis has focused heavily on the nature of the syntrophic intermediates shared between ANME and SRB[10,11,13,34,35]. We currently have an incomplete understanding of the scope of other potential metabolic interdependencies within this long-standing symbiosis. Prior experimental research has demonstrated the potential for nitrogen fixation and exchange in AOM consortia under certain environmental circumstances[12,14,31,78], and in other energy limited anaerobic syntrophies between bacteria and archaea, amino acid auxotrophies are common[79–81]. Comparative analysis of metagenome assembled genomes from several lineages of ANME archaea[15] as well as a subset of syntrophic sulfate-reducing bacterial partners[21] lacked evidence for specific loss of pathways used in amino acid synthesis, and our expanded analysis of SRB here is consistent with these earlier studies. Interestingly, comparative analysis of specific pairings of ANME and their SRB partners revealed the possibility for cobalamin dependency and exchange. Cobamides, also known as the Vitamin B12-type family of cofactors, are critical for many central metabolic pathways[82]. Mechanisms for complete or partial cobamide uptake and remodeling by microorganisms found in diverse environments is common[82]. The importance of exchange of cobamide between gut bacteria, and between bacteria and eukaryotes has been demonstrated[83,84]. In methanotrophic ANME-SRB partnerships, ANME are dependent on cobalamin as a cofactor in their central metabolic pathway and biosynthetic pathways, while Seep-SRB2, Seep-SRB1a, Seep-SRB1g also have essential cobalamin dependent enzymes including ribonucleotide reductase, methionine synthase and acetyl-CoA synthase (**Supplementary Table 8**). This is in contrast with the HotSeep-1 clade, which appear to have fewer cobalamin requiring enzymes and may not have an obligate dependence on vitamin B12. However, HotSeep-1 do possess homologs of BtuBCDF and CobT/CobU, genes that are used in cobamide salvage and remodeling[85] (**Supplementary Table 8**). An absence of cobalamin biosynthesis in either ANME or these three clades of syntrophic SRB would thus necessarily lead to a metabolic dependence on either the partner or external sources of cobalamin in the environment. We observed such a predicted metabolic dependence for Seep-SRB1a within the species Seep-SRB1a sp. 1 (n= 1 genomes), Seep-SRB1a sp. 5 (n=4 genomes), Seep-SRB1a sp. 3 (n=2 genomes), Seep-SRB1a sp. 7 (n=1) and Seep-SRB1a sp. 8 (n=1). All these genomes are missing the anaerobic corrin ring biosynthesis pathway but, some do retain genes involved in lower ligand synthesis (BzaAB)[86] (**Figure 5**). Additionally, recent metatranscriptomic data from an AOM incubation dominated by ANME-2a/Seep-SRB1a associated with Seep-SRB1a sp. 5 (str. SM7059A) that is missing the cobalamin biosynthesis pathways confirmed active expression of cobalamin dependent pathways in the Seep-SRB1a including ribonucleotide reductase and acetyl-coA synthase AcsD[22] suggesting that these syntrophs must acquire cobalamin from their ANME partner or the environment. Interestingly, the predicted cobalamin auxotrophy is not a uniform trait within the Seep SRB1a lineage, with cobamide biosynthesis genes present in the genomes of species Seep-SRB1a sp. 2 (n=3), Seep-SRB1a sp. 4 (n=1), and Seep-SRB1a sp. 9 (n=3). Two of the five Seep-SRB1a species missing cobalamin biosynthesis genes are known to partner ANME-2a while Seep-SRB1a sp. 2 is a known ANME-2c partner as demonstrated by sequencing of single consortia containing ANME-2c and Seep-SRB1a sp. 2 and Seep-SRB1a sp. 4 is sourced from a sample containing ANME-2c and a lower abundance of ANME-2a. The striking absence of cobalamin biosynthesis genes in verified Seep-SRB1a partners of ANME-2a, while the closely related Seep-SRB1a partners of ANME-2c appear to maintain this biosynthetic ability is intriguing and may point to the development of nutritional auxotrophy shaped by the specific syntrophic association between ANME-2a and Seep-SRB1a. Future experimental work will assist with testing this predicted vitamin dependency among the ANME-2a and Seep-SRB1a and other ANME-SRB partner pairings.

**Figure 5.**
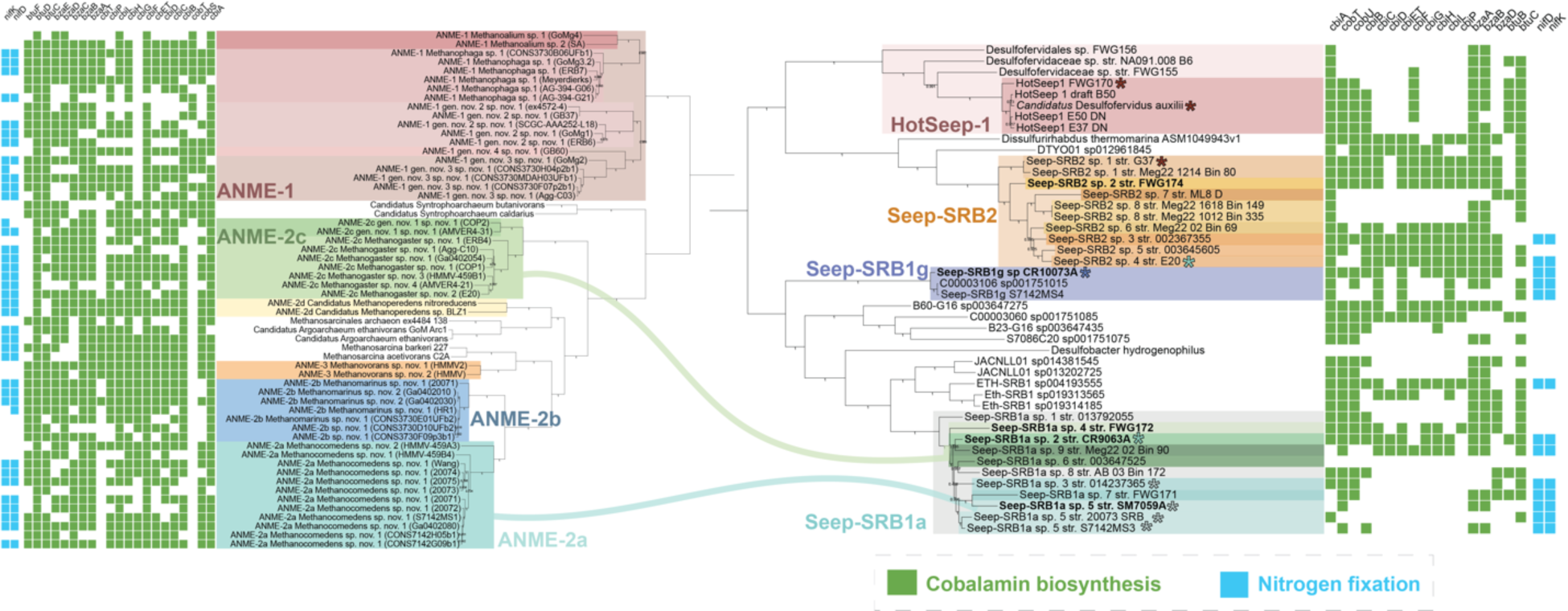
The loss of cobalamin biosynthesis genes in the Seep-SRB1a partners of ANME-2a. On the right, a phylogenetic tree of concatenated ribosomal proteins from all the genomes of syntrophic sulfate reducing bacteria clades – Hot-Seep1, Seep-SRB2, Seep-SRB1g and Seep-SRB1a and related clades, Seep-SRB1c and Eth-SRB1 was made using Anvi’o[1]. On the left, a similar concatenated protein tree was made for ANME genomes highlighting the clades from ANME-1, ANME-2c, ANME-2b and ANME-2a. Lines in green, blue and light teal are used to depict the partnerships between ANME-2c, ANME-2b and ANME-2a. ANME-2c genomes are not separated into those belonging to partners of Seep-SRB2 and Seep-SRB1a. The presence of genes involved in cobalamin biosynthesis and nitrogen fixation are marked in light green and light blue respectively. The proposed type strains are bolded.

The ability to fix nitrogen is found in bacteria and archaea but is relatively rare amongst them[87]. Fixed nitrogen availability can impact the productivity of a given ecosystem. Members of the ANME-2 archaea have been demonstrated to fix nitrogen in consortia [12,13,78] and may serve as a source of fixed nitrogen for methane-based communities in deep-sea seeps[78]. We recently demonstrated that within the ANME-2b/Seep-SRB1g partnership, Seep-SRB1g bacteria can also fix nitrogen[14]. A comparison of the nitrogen fixation ability across ANME and SRB (**Figure 5**), shows that this function is present in the genome representatives of diverse ANME, and also conserved in some syntrophic bacterial partners (Seep-SRB1a and Seep-SRB1g). In the Seep-SRB1a lineage, the nitrogenase operon is retained in both ANME-2a and ANME-2c partners, contrasting the pattern observed with cobalamin synthesis. Interesting, the potential to fix nitrogen occurs in species of Seep-SRB2 that come from psychrophilic deep-sea environments (Seep-SRB2 sp. 4 and Seep-SRB2 sp. 3), while earlier branching clades of Seep-SRB2 adapted to hotter environments (Seep-SRB2 sp. 1 and 2) lack nitrogenases, hinting at potential ecophysiological adaptation to temperature (**Figure 5**). While the ability to fix nitrogen is retained in several clades of syntrophic SRB, previous stable isotope labeling experiments have shown that ANME is the dominant nitrogen fixing partner[12,14,78], a trait consistent with its function as the partner with access to lower potential electrons. Yet, the potential to fix nitrogen is retained in Seep-SRB1a and Seep-SRB1g members (**Figure 5**), and in some cases, have been directly linked to N_2_ fixation in the case of Seep SRB1g[14] or indirectly suggested from the recovery of nifH transcripts belonging to Seep-SRB1a and Seep-SRB1g in seep sediments[13]. These observations indicate that nitrogen sharing dynamics between ANME and SRB is likely more complicated than we have thus far observed and may correspond to differences in environment, or perhaps to specific partnership interactions that require assessment at greater taxonomic resolution.

#### e. Pathways related to biofilm formation and intercellular communication

ANME and SRB form multicellular aggregates in which they are spatially organized in distinct and recognizable ways[30]. ANME-2a/2b/2c and ANME-3 are known to form tight aggregates with their bacterial partners[11,24,26,30]. Members of ANME-1 have been observed in tightly packed consortia with SRB[24], while others some form more loose associations[16,88–90]. In these consortia, archaeal and bacterial cells are often enmeshed in an extracellular polymeric substance[88,91,92]. In large carbonate associated mats of ANME-2c and ANME-1 and SRB from the Black Sea, extractions of exopolymers consisted of 10 % neutral sugars, 27% protein and 2.3% uronic acids[91]. This composition is consistent with the roles played by mixed protein and extracellular polysaccharide networks shown to be important for the formation of conductive biofilms in *Geobacter sulfurreducens*[93], the formation of multicellular fruiting bodies from *Myxococcus xanthus*[94–96] and the formation of single-species[97] and polymicrobial biofilms[98]. Important and conserved features across these biofilms are structural components made up of polysaccharides, cellular extensions such as type IV pili and matrix binding proteins such as fibronectin containing domains[99]. Functional components of the biofilm matrix such as virulence factors in pathogens[100] and extracellular electron transfer components[93] are variable and depend on the lifestyle of the microorganism. Mechanisms to sense or recognize other cells is another common adaptation to growth in a biofilm. For e.g., in Vibrio biofilms the secreted protein RbmA is known to be important for kin selection and the prevention of other microorganisms from invading Vibrio biofilms[101]. Guided by the molecular understanding of mechanisms and physiological adaptation to microbial growth in biofilms, we examined the genomic evidence for similar adaptations in the syntrophic SRB in consortia with ANME archaea, focusing on structural and functional components of biofilms as well as proteins implicated in partner identification (**Figure 6**).

**Figure 6.**
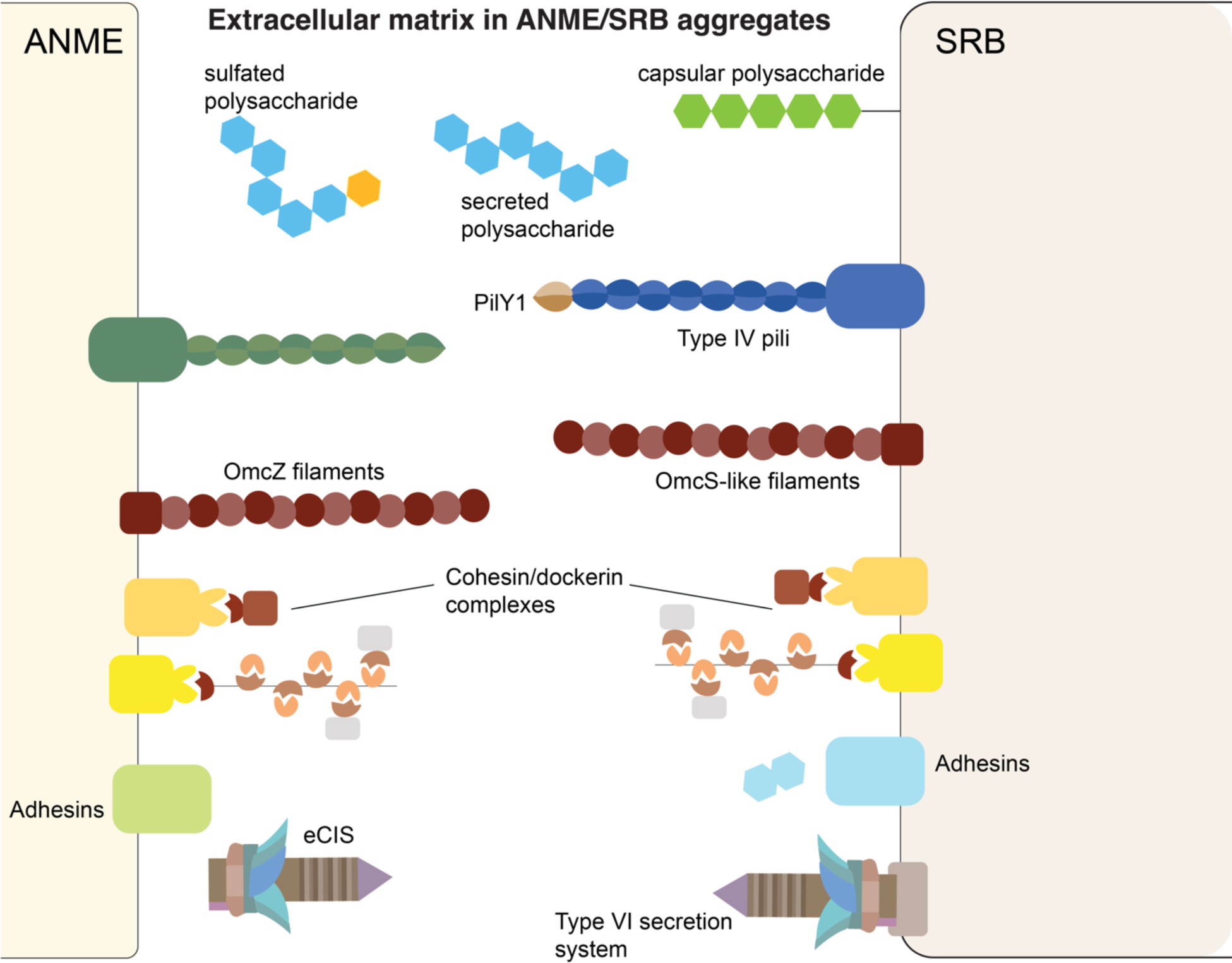
Putative physiological factors involved in ANME/SRB aggregate formation. Extracellular polysaccharides and protein complexes implicated in the formation of the extracellular matrix in ANME-SRB aggregates are visualized as cell-surface embedded or secreted. The capacity for biosynthesis of sulfated polysaccharides is present in three of the syntrophic SRB clades – Seep-SRB2, Seep-SRB1a and Seep-SRB1g. Type VI secretion systems and extracellular contractile injection systems (eCIS) are likely important for intercellular communication between ANME and SRB.

Our analysis of syntrophic SRB genomes showed the presence of multiple putative polysaccharide biosynthesis pathways in different SRB lineages including secreted extracellular polysaccharide biosynthesis pathways and capsular polysaccharide biosynthesis pathways (**Supplementary Table 9**). In particular, homologs of the pel biosynthesis pathway (PelA, PelE, PelF and PelG), first identified in *Pseudomonas aeruginosa*[102,103] were present in almost all Seep-SRB1g and Seep-SRB1a genomes (**Supplementary Figures 12, 13**). These homologs are part of a conserved operon in these genomes which includes a transmembrane protein that could perform the same function as PelD, which along with PelE, PelF and PelG forms the synthase component of the biosynthetic pathway and enables transport of the polysaccharide pel across the inner membrane[103]. Metatranscriptomic data confirms this operon is expressed and was significantly downregulated when methane-oxidation by ANME-2a was decoupled from its syntrophic Seep SRB1a partner with the addition of AQDS[22]. This biosynthesis pathway is absent in the nearest evolutionary neighbors of Seep-SRB1a and Seep-SRB1g, Eth-SRB1 and Seep-SRB1c respectively, suggesting that the presence of the pel operon could serve as a better genomic marker for syntrophic interaction with ANME-2a, ANME-2b and ANME-2c than the presence of the oetB-type conduit. The pel operon was also detected in one of the Seep-SRB2 genomes but, is not conserved across this clade. In Seep-SRB2 clades, multiple capsular polysaccharide biosynthesis pathways are conserved. This includes a neuraminic acid biosynthesis pathway, a sialic acid capsular polysaccharide widely associated with intestinal mucous glycans and used by pathogenic and commensal bacteria to evade the host immune system[104] (**Supplementary Figure 14**). These differences in polysaccharide biosynthesis pathways are likely reflected in the nature of the EPS matrix within each ANME-SRB aggregate.

Members of the thermophilic HotSeep-1 syntrophic SRB also encode for multiple putative polysaccharide biosynthesis pathways, including a pathway similar to the xap pathway in *G. sulfurreducens* (**Supplementary Figure 15**). The role of polysaccharides in the formation of conductive extracellular matrices and in intercellular communication is just beginning to be understood but, they appear to be essential to its formation. For example, the mutation of the xap polysaccharide biosynthesis pathway in *G. sulfurreducens* eliminated the ability of this electrogenic bacteria to reduce Fe (III) reduction in the bacterium[93] and affected the localization of key multiheme cytochromes *c* OmcS and OmcZ and structure of the biofilm matrix[105] suggesting that the EPS matrix contributes a structural scaffold for the localization of the multiheme cytochromes. Similarly, the cationic polysaccharide pel in *P. aeruginosa* biofilms has recently been shown to play a role in binding extracellular DNA or other anionic substrates together forming tight electrostatic networks that provide strength to the extracellular matrix[106] and may offer a similar role in Seep SRB1a and 1g consortia. Based on the reported chemical composition of EPS from the Black Sea ANME-SRB biofilm[91], alongside TEM compatible staining of cytochromes *c* in the extracellular space between ANME and SRB[10,11,24], and the genomic evidence provided here of conserved polysaccharide biosynthesis pathways point to the existence of a conductive extracellular matrix within ANME-SRB consortia that has features similar to *Geobacter* biofilms[93]. While these conductive biofilms are correlated with the presence of secreted polysaccharides, the highly conserved capsular polysaccharides common in Seep SRB2 likely play a different role. In *Myxococcus xanthus*, the deletion of capsular polysaccharides leads to a disruption in the formation of multicellular fruiting bodies, suggesting a possible role for capsular polysaccharides in intercellular communication[107]. This is consistent with the universal role of O-antigen ligated lipopolysaccharides in cell recognition and the Seep SRB2 capsular polysaccharides may serve a similar purpose in consortia with ANME archaea, either influencing within population interactions, or potentially mediating kin recognition.

In addition to polysaccharides, type IV pili are well-conserved in the syntrophic SRB genomes (**data not shown**). Consistent with this, HotSeep-1 pili are known to be well-expressed[10] and were shown to be non-conductive and proposed to play a role in intercellular communication[108]. PilY1 is a subunit of type IV pili that is known to facilitate to promote surface adhesion in *Pseudomonas* and intercellular communication in multi-species *Pseudomonas* biofilms[109]. Other adhesion related proteins in the syntrophic SRB include cohesin and dockerin domain containing proteins such as those previously identified in ANME[15], immunoglobulin-like domains, cell-adhesin related domain (CARDB) domains, bacterial S8 protease domains, PEB3 adhesin domains, cadherin, integrin domains and fibronectin domains (**Figure 6, Supplementary Table 10**). Fibronectin domains are found in the one of the cytochromes *c*, oetF that is likely part of the extracellular electron transfer conduit. This domain might interact with the conductive biofilm matrix itself or serve as a partnership recognition site. Our analysis of the SRB adhesins suggests that some adhesins are conserved across a given syntrophic clade, while others appear to be more species or partnership specific. For e.g., while PilY1 is conserved across Seep-SRB2, the cohesin/dockerin complexes that are conserved in Hot-Seep1 and Seep-SRB1g are thus far found only in Seep-SRB2 sp. 4 and 8. Analysis of gene expression data suggest that in the Hot-Seep1/ANME-1 partnership, PilY1, an adhesin with an immunoglobulin-like domain and adjacent cohesin/dockerin domains might play a role in the syntrophic lifestyle[24]. In the ANME-2c/Seep-SRB2 partnership, PilY1, cohesin/dockerin complexes and a protein with a CARDB domain are highly expressed[24]. Curiously, in the Seep-SRB2 partnering with ANME-1, we could only identify one moderately expressed adhesin with a fibronectin domain[24] (**Supplementary Table 10**). Interestingly, we note the presence and high levels of expression of cohesin/dockerin domains in both ANME-2c and their verified Seep-SRB2 partner[24], and the presence of fibronectin domains in both ANME-2a and their Seep-SRB1a partner **Supplementary Table 10**) suggesting that perhaps both partners within a partnership express and secrete similar kinds of extracellular proteins. This might serve as a mechanism for partnership sensing, While our analysis and that of earlier research into adhesins present in ANME[15] identify a number of conserved and expressed adhesins, further work is needed to investigate their potential role in aggregate formation.

Extracellular contractile injection systems (eCIS) that resemble phage-like translocation systems (PLTS) are found in some syntrophic SRB genomes (**Supplementary Table 11**, **Supplementary Figure 16**) although they are not as widely distributed as in ANME[15]. Typically, the eCIS bind to a target microorganism and release effector proteins into its cytoplasm. eCIS have been shown to induce death in worm larvae, induce maturation in marine tubeworm larvae[110] and found to mediate interactions between the amoeba symbiont and its host[111]. In ANME-SRB consortia, they might play a similar role with ANME releasing an effector protein into SRB, perhaps an effector molecule to promote the formation of a conductive biofilm or adhesins. Type VI secretion systems (T6SS) are similar to eCIS in facilitating intercellular communication between microorganisms. However, the primary distinction between them is that T6SS are membrane-bound while eCIS appear to be secreted to the extracellular space[112,113]. Interestingly, T6SS appear to be present in the ANME-2a partner Seep-SRB1a but absent in the ANME-2c partner Seep-SRB1a suggesting that they might play a role in mediating partnership specificity. Our analysis identified many conserved mechanisms for biofilm formation and intercellular communication in SRB to complement the pathways previously identified in ANME. Significantly, several polysaccharide biosynthesis pathways and adhesins were absent in the closest evolutionary neighbors of SRB indicating that adaptation to a syntrophic partnership with ANME required not just metabolic specialization but adaptation to a multicellular and syntrophic lifestyle.

### Physiological adaptation of syntrophic SRB to partnerships with ANME

To better understand the evolutionary adaptations acquired by syntrophic SRB to form partnerships with ANME, we mapped the presence and absence of the above-mentioned pathways in central metabolism, nutrient sharing, biofilm formation, cell adhesion and partner identification across each of the syntrophic SRB clades and their nearest evolutionary neighbors from the same bacterial order (**Supplementary Table 12**, **Supplementary Table 13**). For e.g., the presence of the extracellular electron transfer conduit OetABI in the Seep-SRB1a clade is nearly universal but, this trait is absent in the Desulfobacterales order that Seep-SRB1a belongs to, suggesting strongly that this machinery was horizontally acquired possibly in Seep-SRB1a or a closely related ancestor within the same family that includes Eth-SRB1. In contrast, most genomes in the order that Seep-SRB1g belongs to contain hydrogenases. However, hydrogenases are lacking in the syntrophic clade Seep-SRB1g implying that this trait was lost in the process of specialization to a partnership with ANME-2b. In addition to inferring adaptation based on presence and absence, phylogenetic trees were generated for at least one representative gene from each identified characteristic to corroborate the possibility of horizontal gene transfers. These trees provide further insight into the adaptation of various traits, the likely source of the genes received horizontally and in the case of Hot-Seep1 and Seep-SRB2 sp. 1 demonstrate the transfer of OetABI from one syntrophic clade to another. With the trees, we were able to also identify those genes that were vertically acquired but heavily adapted for the respiratory pathways receiving DIET electrons, for example Tmc (**Supplementary Figure 8**). A brief summary of the gene gains and losses is provided in **Figure 7** and **Supplementary Table 13**. Our analysis suggests that some traits are associated with partnerships with different ANME. The pel operon present in Seep-SRB1g and Seep-SRB1a is more closely associated with aggregates formed with the ANME-2a/b/c species rather than ANME-1. Similarly, the capsular polysaccharide pseudaminic acid is present in those species of Seep-SRB1a that are associated with ANME-2c but absent in those species partnering ANME-2a suggesting that this polysaccharide might play a role in partnership identification and aggregate formation. Curiously, many of the adhesins we identified in the syntrophic SRB genomes have few close homologs in the NCBI nr dataset and almost no homologs in the nearest evolutionary neighbors (**Supplementary Table 13**), indicating that these proteins are likely highly divergent from their nearest ancestors. This is consistent with faster adaptive rates observed in extracellular proteins[114] and in general, the higher horizontal gene transfer rates for cell-surface proteins[115].

**Figure 7.**
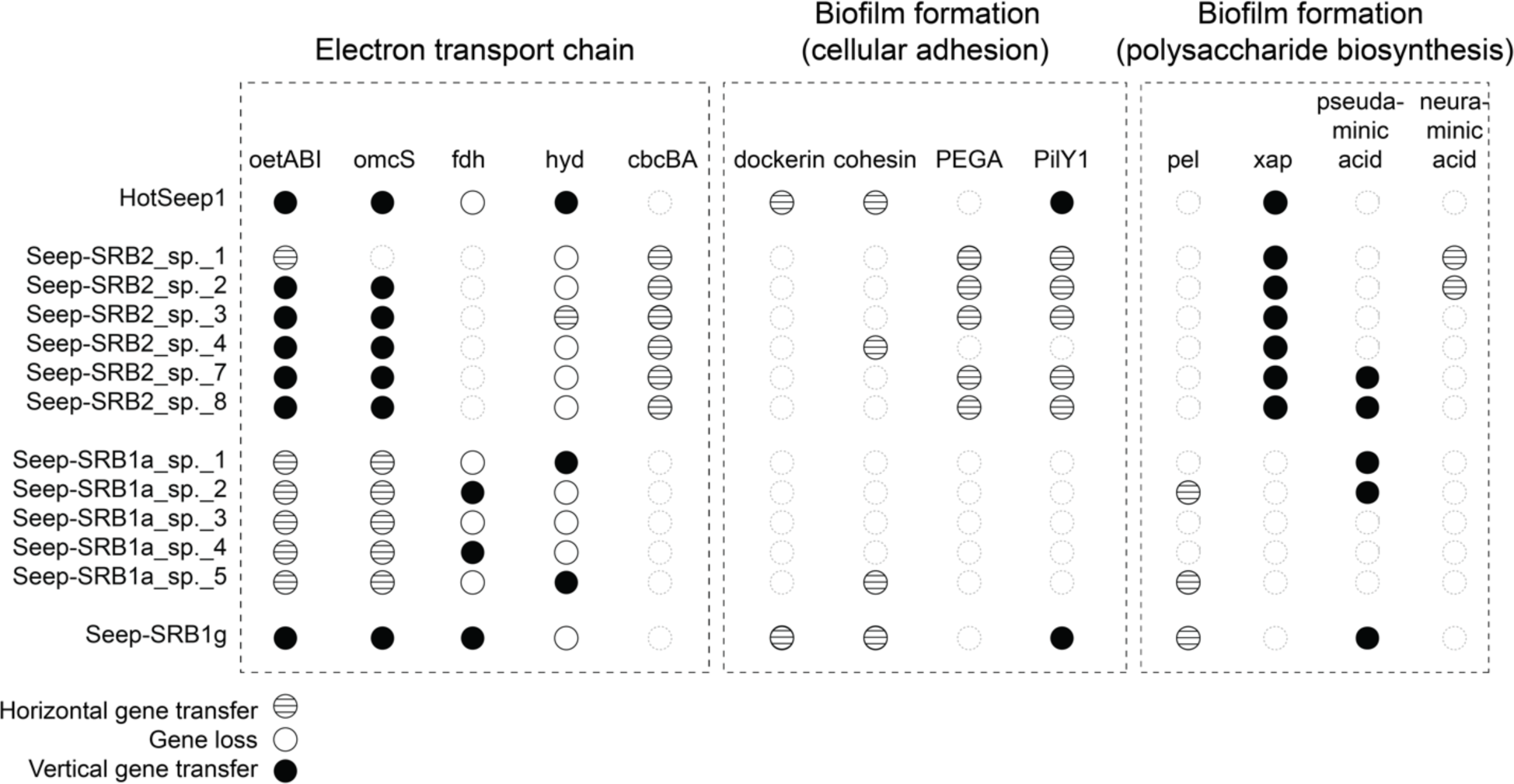
A summary of important gene loss and gain events in the physiological adaptation of sulfate reducing bacteria that led to a syntrophic partnership with ANME. The presence and absence of genes involved in the electron transport chain, nutrient sharing, biofilm formation and cellular adhesion are listed in **Supplementary Table 12**. We identified genes that were potentially gained, lost, or biochemically adapted using a comparative analysis of the presence a given gene in a syntrophic clade in its order-level taxonomic background. For e.g., if a gene is present in a syntrophic SRB clade and is present in fewer than 30 % of the remaining species in a given order, this gene is considered a likely horizontally transferred gene. The likelihood of horizontal transfer is then further corroborated with a phylogenetic tree of that gene generated with close homologs from NCBI and our curated dataset. The secondary analysis of the likelihood of gene gains and losses is present in **Supplementary Table 13**.

With our analysis, we identified many genes and traits that are correlated with a syntrophic partnership with ANME, but it is less easy to identify whether there are essential traits were for the formation of this syntrophic partnership. The complete conservation of the OetB-type or other EET cluster (such as OmcKL) suggests these are essential, but not sufficient, for the formation of this partnership since the multi-heme cytochrome conduits themselves are present in many organisms not forming a syntrophic partnership with ANME. There is also a strong signature for the presence of a secreted polysaccharide pathway such as the pel operon in Seep-SRB1a and Seep-SRB1g and a xap-like polysaccharide in Hot-Seep1 and Seep-SRB2. With these components, a conductive biofilm matrix can be established, but the means of partnership recognition and communication between the archaea and bacteria are less clear. As suggested previously[15], the near complete conservation of the extracellular contractile injection systems (eCIS) in ANME might play a role in partnership identification. The target receptor of the eCIS is unclear but the presence of conserved capsular polysaccharides in SRB that often are the target of bacteriophages and pathogens is suggestive as a possible site for binding. Likewise, the high levels of expression of cohesin and dockerin complexes by both ANME and SRB in the ANME-2c/Seep-SRB2 partnership is indicative of a role in syntrophic partnership[24]. In Seep-SRB1a there are conserved fibronectin domains that likely bind the biofilm matrix and Seep-SRB2 has a conserved cell-surface protein with a PEGA sequence motif (**Supplementary Table 12**). The respiratory pathways for DIET are present ancestrally in Desulfofervidales and the order C00003060, suggesting that the syntrophic partners in these orders, Hot-Seep1 and Seep-SRB1g respectively acquired the pathways needed for aggregate formation (such as adhesins, the pel polysaccharide biosynthesis pathway) after. In contrast, both Seep-SRB2 and Seep-SRB1a contain respiratory traits (CbcBA and OetABI respectively) that were likely absent in their ancestors (**Figures 2, 4**). This indicates that more steps were required for the adaptation of these clades to a syntrophic partnership with ANME. The greater diversity within these clades may also reflect the co-diversification that must have occurred in these clades that are known to partner multiple ANME. However, there is insufficient evidence to rule out the possibility of promiscuous partnership formation with multiple ANME within each SRB clade. In these cases, the observed diversity must be driven by other factors. Within each syntrophic clade, the complete conservation of respiratory pathways and some polysaccharide biosynthesis pathways suggests that these traits were acquired early within the evolutionary trajectory of these organisms and stabilized within their genomes. The adaptation towards extracellular electron transfer and the formation of conductive biofilms was likely driven by a greater selection pressure than the adaptation to a specific ANME partner. Hence, the gain and loss of specific adhesin and matrix binding proteins is more dynamic. If this were the sequence in which syntrophic SRB diversified and adapted to a partnership with ANME, it would be consistent with the evolution of such a partnership occurring in an early environment where metal cycling was dominant with both the ANME and SRB already capable of extracellular electron transfer to metals.

Another aspect of the adaptation of syntrophic SRB is the high number of inter-clade transfers. In addition to the likely transfer of OetABI (**Supplementary Figure 7**), we also note a high degree of similarity between the proteins of the following components in different clades of syntrophic SRB - cohesin/dockerin modules, the OmcKL conduit, and enzymes in the pel and xap polysaccharide biosynthesis pathways. These appear to be the result of inter-clade transfers and the high number of transfers might imply that a mechanism promoting the exchange of DNA exists in this environment between ANME and SRB, either through a viral conduit or perhaps with the eCIS carrying DNA as cargo. Further analysis is needed to identify the number of transfer and the sources of transfers. In fact, a thorough accounting of these horizontal gene transfers combined with molecular clock dating might provide insight into the timeline and the relative age of the different ANME/SRB partnerships. Our phylogenomic analysis places the verified ANME-2c partners as ancestral to the ANME-2a partners within the Seep-SRB1a clade (**Figure 1**, **Supplementary Figure 1**). Within the Seep-SRB2 clade, the topology places an ANME-1 partner as basal to the remaining Seep-SRB2 and the only verified ANME-2c partner as one of the later branching members (**Figure 1**, **Supplementary Figure 1**). Earlier research places ANME-1 as the deepest branching lineage of ANME[15] and this relative ancestry of partners might suggest that Seep-SRB2 is older than Seep-SRB1a. However, it appears that ANME-1 acquired its mcr through horizontal gene transfer[15], and we have insufficient data to know when this occurred. Thus, we cannot know that ANME-1 was methanotrophic when it diverged from the Methanomicrobiales. These observations suggest that we cannot constrain the emergence of AOM solely through the relative branching patterns of the various ANME and SRB clades. A more thorough reconstruction of the adaptive gene transfers using the framework established for ANME and in this work for syntrophic SRB would provide insight into the evolution of this biogeochemically important syntrophic partnership.

## Conclusions

This comparative genomic analysis of the major ANME-partnering SRB clades provides a valuable metabolic and evolutionary framework to understand the differences between the various syntrophic sulfate reducing partners of anaerobic methanotrophic archaea and develop insight into their metabolic adaptation. In this work we show that the electron transport chains of the different syntrophic SRB partners of ANME are adapted to incorporate extracellular electron transfer conduits that are needed for direct interspecies electron transfer. Groups including the Seep-SRB2 appear to have acquired cytoplasmic membrane complexes that can function with the EET conduits, while Seep-SRB1a clades have adapted existing inner-membrane complexes for interaction with the EET conduit. Electron bifurcation also appears to be common across the syntrophic lineages and is often coupled to the cytoplasmic machinery, and likely provides an advantage in low energy environments. We also show that the co-evolution between different ANME and SRB partners may have resulted in nutritional interdependencies, with cobalamin auxotrophy observed in at least one of the specific syntrophic SRB subclades. Our genome-based observations provide insight into the various adaptations that are correlated with the formation of different ANME-SRB partnerships. These adaptive traits appear to be related with mechanisms driving other ecological phenomena such as biofilm formation and non-obligate syntrophic interactions. The identification of these traits allowed us to posit important steps in the evolutionary trajectory of these sulfate reducing bacteria to a syntrophic lifestyle. While the full import of these observations is not yet clear, they offer a roadmap for targeted physiological investigations, and phylogenetic studies in the future.

## Acknowledgment

We thank the DOE and the Moore Foundation for funding this research (Principle Investigator: Dr. Victoria J. Orphan).We acknowledge the Dalio Foundation and Woods Hole Oceanographic Institute for supporting the NA091 research cruise to South Pescadero Basin on E/V Nautilus operated by the Ocean Exploration Trust in October-November 2017. The work (Award doi: 10.46936/fics.proj.2017.49956/60006219) conducted by the U.S. Department of Energy Joint Genome Institute (https://ror.org/04xm1d337), a DOE Office of Science User Facility, is supported by the Office of Science of the U.S. Department of Energy operated under Contract No. DE-AC02-05CH11231. We would also like to thank Magdalena Mayr for her thoughtful comments on this manuscript and Fernanda Jimenez-Otero for sharing her thoughts and expertise in the field of extracellular electron transfer. We are also grateful to Alon Philosof, Aditi Narayan, Kriti Sharma and James Hemp for many productive discussions on broad scientific questions in microbial ecology and evolution, metabolism and scientific writing that lent itself to the framing of this manuscript.

## Materials and Methods

### Sampling locations and processing of samples

Push-core samples of seafloor sediment were collected from different locations on the Costa Rica margin during the AT37-13 cruise in May 2017 (sample serial numbers: #10073, #9063), southern Pescadero Basin[1] during the FK181031 cruise on R/V Falkor operated by the Schmidt Ocean Institute in November 2018 (sample serial number: #PB10259, live incubation of the top 3 cm section of push core #FK181031-S193-PC3)[2] and from Santa Monica Basin during the in May 2013 (sample serial numbers: #7059). Sediment push-cores retrieved from the seafloor were sectioned into 1-3 cm sediment horizons. At the time of shipboard processing, ∼2mL of the sediment was sampled for DNA extraction and FISH analysis and the rest was saved in Mylar bags under an N_2_ atmosphere at 4 ℃ for sediment microcosm incubations. Microbial mat sample #14434 was collected from Santa Monica Basin during the WF02-20 cruises in February 2020. Rock samples were retrieved from South Pescadero Basin[2] during the NA091 cruise on E/V Nautilus operated by the Ocean Exploration Trust in October-November 2017 and the FK181031 cruises (sample serial numbers: NA091-R045, NA091-R008 and, #12019, #11946, #11719, respectively) and saved in Mylar bags under an N_2_ atmosphere at 4 ℃. Further details on sampling locations are available in Supplementary Information.

Sediment horizons from samples 10073, 9063 and 7059 were incubated in artificial sea water as previously described[3,4] with CH_4_ and 250 µM L-Homopropargylglycine (HPG) at 4 ℃. Once the presence of metabolically active ANME-SRB in these microcosms was confirmed by the accumulation of sulfide combined with observation of incorporation of HPG by BioOrthogonal Non-Canonical Amino Acid Tagging (BONCAT), samples from these incubations were used for sorting of single-aggregates by BONCAT-FACS as described below. #FK181031-S193-PC3 was incubated in anaerobic artificial sea water without electron donor at 24°C. Rock #NA091-R045 was incubated in anaerobic artificial sea water supplemented with pyruvate at 24°C. Rock samples from S. Pescadero basin (#11946, #11719 and #12019) were also incubated with artificial sea water, and CH_4_ at 50 ℃.

### DNA extraction followed by metagenome sequencing for samples #11946, #11719, #12019, #NA091-R045, #NA091-R008, #PB10259 and #14434

For incubations of carbonate samples #11946, #11719 and #12019, DNA was extracted from approximately 500 mg of crushed rock samples using a modified version of the Zhou protocol[5] as follows. Prior to the incubation with proteinase K, the sample was incubated with lysozyme (10 mg ml-1) for 30 min at 37 °C; 10% SDS was used for incubation; after SDS incubations, the sample was extracted twice by adding 1 volume (1 mL) of phenol/chloroform/isoamylalcohol (25:24:1) with incubation for 20 min at 65 °C followed by centrifugation; in the final step, the DNA was eluted in 40 µL of TE 1x buffer. 250 mg of sediment sample #PB10259 and microbial mat sample #14434 were extracted using the QIAGEN Power Soil Kit. 500 mg of crushed carbonate samples #NA091-R045, #NA091-R008 were also extracted using the QIAGEN Power Soil Kit.

For samples #PB10259, #14434, NA091-R045, DNA libraries were prepared using the NEBNext Ultra kit and sequenced at Novogene with the instrument HiSeq4000. A Library was also prepared using the NEBNext Ultra kit for NA091-008. This sample was sequenced at Quick Biology (Pasadena, CA) with a HiSeq2000 using a 2×150 protocol. DNA libraries for samples #11946, #11719 and #12019 were prepared using the Nextera Flex kit and also sequenced at Novogene on the HiSeq4000. After sequencing of NA091-R45, primers and adapters were removed from all libraries using bbduk[6] with mink=6 and hdist=1 as trimming parameters, and establishing a minimum quality value of 20 and a minimal length of 50 bp.

### Assembly and binning of metagenomes from samples samples #NA091-R045, #NA091-R008, #PB10259, #14434, #11946, #11719 and #12019

Metagenomes from samples #PB10259, #14434, #11946, #11719 and #12019 were assembled individually using SPAdes[7] v3.14.1, and each resulting assembly was binned using metabat v2.15[8]. Automatic prediction of function for genes within the various MAGs was performed using prokka v.1.14.6[9]. The reads of the DNA libraries derived from the rock sample (NA091-R008) were assembled individually using SPAdes v.3.12.0. From the de-novo assemblies for NA091-R008, we performed manual binning using Anvio v.6[10]. We assessed the quality and taxonomy affiliation from the obtained bins using CheckM[11] and GTDB-tk[12]. Genomes of interest affiliated to Desulfobacterota were further refined via a targeted-reassembly pipeline. The trimmed reads for the NA091-008 assembly were mapped to the bin of interest using bbmap[6] (minimal identity of 0.97), then the mapped reads were assembled using SPAdes and finally the resulting assembly was filtered discarding contigs below 1500 bp. This procedure was repeated for 13-20 cycles for each bin, until the bin quality did not improve any further. Bin quality was assessed based on the completeness, contamination (< 5%), N50 value and number of scaffolds of the bin using checkM. The resulting bins were considered as metagenome-assembled genomes (MAGs). Automatic prediction of function for genes within the various MAGs was performed using prokka v.1.14.6[9] and curated with the identification of Pfam[13] and TIGRFAM[14] profiles using HMMER v.3.3[15]; KEGG domains[16] with Kofam[17] and of COGs and arCOGs motifs[18] using COGsoft[19].

### Fluorescent-sorting of metabolically active single aggregates from samples #10073, #9063 and #7059 followed by sequencing

Sediment-extracted consortia from samples #10073, #9063 and #7059 were analyzed. Individual ANME:SRB consortia were identified and sorted using fluorescent signal, as previously described[4]. The SYBR-Green dye was excited using a 488-nm laser, and fluorescence was captured with a 531-nm/ 30-nm filter. Gates were defined using a forward scatter (FSC) vs. 531-nm emission plot, and events with a fluorescent signal brighter than >90% of aggregates in the negative control were captured. For sample #10073, 50 consortia were sorted into 1.5 mL tubes and stored at 4 C for sequencing. For samples #9063 and #7059, 28 and 19 consortia were sorted respectively.

Single consortia were lysed and DNA was amplified using multiple displacement amplification (MDA) protocol as previously described[20]. The amplified DNA was sheared, attached to Illumina adapters, and sequenced using the Illumina NextSeq-HO method. Only metagenomes from two sorted aggregates from each of the samples# 10073, #9063 and #7059 were used in this study.

### Assembly and binning of single aggregate metagenomes from samples #10073, #9063 and #7059

Metagenomes were assembled using SPAdes spades v. 3.13.0 and annotated using the Integrated Microbial genomes (IMG) annotation pipeline. As the single aggregate metagenomes represent extremely reduced communities, and the MDA precludes traditional contig binning by coverage, the metagenomes were binned using a manual approach based on sequence composition and taxonomic assignment of the genes. Manual binning was performed using a principal component analysis (PCA) of the tetramer frequency of the contigs, calculated using calc.kmerfreq.pl(https://github.com/MadsAlbertsen/miscperlscripts/). Taxonomic affiliation of the genes on the contigs was taken from the IMG annotation, and the percentage of genes on each contig annotated as “Archaeal” or “Bacteria” was used to corroborate the clustering in the PCA plot. Jupyter notebooks used for the binning are available at https://github.com/dspeth. Automatic metabolic prediction of the MAGs was performed using prokka v.1.14.6[9].

### Taxonomic classification of metagenome bins from various syntrophic sulfate reducing bacteria

Single copy marker genes identified in the “Bacteria 71” gene set included in Anvio[10] were extracted from each of the syntrophic SRB genomes and all genomes within the phylum Desulfobacterota available in release89 of the Genome Taxonomy Database[21]. A concatenated gene alignment was generated using MUSCLE[22] as part of the anvio script “anvi-get-sequences-for-hmm-hits”. A phylogenetic tree was inferred using FastTree as per the Anvio-7 pipeline using the command “anvi-gen-phylogenomic-tree”, in order to provide a phylogenetic context for each of the four SRB clades. We corroborated our phylogenetic placement with the classification provided by GTDB-tk[12]. Additionally, we assessed the extent of taxonomic diversity within the four clades by calculating the average nucleotide identity (ANI) and 16S rRNA sequence similarity between different organisms that belong to each clade using PyANI[23] in Anvio-7[10]. An 95 % ANI value of 95%[24] and 98.65 % similarity in 16S rRNA[25] were used as cut-offs to delineate different species.

### Phylogenetic analysis of OetI, the outer-membrane beta barrel forming protein in the HmlB-type cluster

OetI here refers specifically to the outer-membrane beta barrel forming protein in the Oet-type cluster implicated in direct interspecies electron transfer between ANME and SRB. All OetI sequences were identified in the genomes of the syntrophic SRB clades by using BLASTP[26] with the query OEU57520.1 from Seep-SRB1g sp. C00003106 and an e-value of e-30. When no OetI hits were found in the syntrophic SRB genomes using this query, we tested for the existence of a beta barrel within ten genes of every multiheme cytochrome that contained more than 5 heme *c* binding motifs using PRED-TMBB[27]. In this way, we identified seventeen EET gene clusters in the syntrophic SRB genomes and five clusters from non-syntrophic Seep-SRB1c, Desulfofervidales and Dissufuribacterales. Protein sequences of OetI from each of these clusters were used as queries to extract all the closest homologs for each of these OetI sequences from the NCBI database. This search was performed using BLASTP with an e-value cut of 1e-5. The extracted sequences were aligned and manually curated to eliminate sequences that were too short and to remove non-specific hits. A phylogenetic tree was inferred using IQ-TREE2[28], a Dayhoff model of substitution and 1000 ultrafast bootstrap iterations and visualized using the iTOL web server[29].

### Phylogenetic analysis of other respiratory proteins

All sequence alignments used for analysis of respiratory proteins were made using MUSCLE[22], and visualized using Jalview[30]. Phylogenetic trees of all proteins were inferred using IQ-TREE2[28] except for the following - OetB, omcX, TmcD and TmcA. Phylogenetic trees for OetB, omcX, TmcD and TmcA were inferred using RAxML[31]. RAxML trees were inferred using a Dayhoff model of rate substitution and 100 bootstraps. IQ-TREE trees were inferred using 1000 ultrafast bootstraps while the models were automatically selected by IQ-TREE using the Bayesian Information Criterion (BIC). The models used for each specific tree are available in **Supplementary Table 14**.

### Sequences analysis of all cytochromes *c*

All cytochromes *c* were identified from the MAGs of syntrophic SRB by employing a word-search method with a custom python script by querying for the commonly found ‘CxxCH’ motif in cytochromes *c*. Once these sequences were extracted, they were aligned using MUSCLE[22]. Clusters were identified depending on the presence of well-defined regions using visual inspection. The clusters were then tabulated in **Supplementary Table 6**. The cellular localization of cytochromes *c* was inferred either from the cellular localization of homologous cytochromes *c* from Desulfuromonadales, or using Signal P-5.0[32].

### Sequences analysis of all putative adhesins

Adhesins were identified from the MAGs of syntrophic SRB by using the ‘all-domain’ annotation feature on KBase as previously described[33,34]. Once putative domains were predicted, we extracted the coding features that corresponded to all putative adhesins based on searches for the words ’integrin’, ’adhesin’, ’cohesin’, ’dockerin’, ’fibronectin’, ’PilY’ and ’immunoglobulin in the domain descriptions. The proteins corresponding to these results were aligned them using MUSCLE[22]. Adhesin clusters were identified depending on the presence of well-defined regions using visual inspection and then additionally verified by use of the NCBI Conserved Domain database[35]. The adhesins were then tabulated in **Supplementary Table 10**.

### Identification of putative polysaccharide biosynthesis pathways

Once, the syntrophic SRB genomes were annotated using the Prokaryotic Genome Annotation Pipeline (PGAP)[36], we identified polysaccharide biosynthesis pathways by looking for the presence of glycosyl transferases, aminotransferases, sugar transporters and polysaccharide biosynthesis proteins. If gene cassette structures followed known operon structures of ABC transporter-type, Wzx/Wzy or synthase type pathways[37], they were retained and tabulated in **Supplementary Table 9** and visualized in **Supplementary Figures 12-15.**

Evolutionary analysis of important genes to identify gains, loss and biochemical adaptation We tabulated the presence and absence of 33 traits that we propose are important for the formation of ANME-SRB partnership in each syntrophic SRB clade and the taxonomic order from which they originate. The presence and absence was identified using BLASTP searches with a query sequence or HMM as listed in **Supplementary Table 13**. If a gene is present in over 30 % of non-syntrophic relatives in a given order, it is considered as present in this order or in a syntrophic SRB clade, it is consider present in this taxonomic clade. If a gene is present in the order and in the syntrophic SRB clade that belongs to this order, the gene is considered to be vertically transferred. If a gene is present in the order but, absent in the syntrophic SRB, the gene is considered to be lost. If a gene is present in the syntrophic SRB but absent in the order it belongs to, the gene is considered to be horizontally acquired. The last assumption is corroborated as much as possible by gene trees deposited in Supplementary Information.

### Data availability for metagenome sequences

Metagenome assemblies from sorted aggregates that we used here are made available on the IMG database under the following IDs – 3300036218, 3300036221, 3300036226, 3300036304, 3300036329, 3300036259. The 14 metagenome assembled genomes (MAGs) of syntrophic SRB from this paper are available on NCBI under the BioProject PRJNA762493.

