## Supplementary Figures for "Physiological adaptation of sulfate reducing bacteria in syntrophic partnership with anaerobic methanotrophic archaea"

### *Table of Contents*


| | |
|---|---|
| Supplementary Figure 1**. Taxonomic separation of syntrophic sulfate reducing bacteria using average nucleotide identity** | **2** |
| **Supplementary Figure** 2**. Comparison of 16S rRNA and 23S rRNA phylogeny of organisms from the phylum Desulfobacterota.** | **3** |
| **Supplementary Figure** 3**. Phylogeny of RpoB from organisms within the phylum Desulfobacterota.** | **4** |
| **Supplementary Figure 4. Placement of various Seep-SRB1 clades within the phylum Desulfobacterota.** | **5** |
| **Supplementary Figure 5. Pan-genome analysis of Seep-SRB1a metagenomes.** | **6** |
| **Supplementary Figure 6. Pan-genome analysis of Seep-SRB2 metagenomes.** | **7** |
| Supplementary Figure 7**. Phylogenetic placement of the outer membrane beta barrel,** OetI **from the putative DIET cluster.** | **8** |
| Supplementary Figure 8**. The Tmc complex in Seep-SRB1a and Seep-SRB1g are divergent.** | **9** |
| **Supplementary Figure 9. Distribution of QrcABCD and RnfABCDEG in Desulfobacterota.** | **10** |
| **Supplementary Figure 10. Gene neighborhoods of various HdrA containing complexes and carbon fixation pathways in syntrophic SRB.** | **11** |
| **Supplementary Figure 11. Operons of Flx-Hdr complexes and gene neighborhood of putative formate utilizing proteins.** | **12** |
| **Supplementary Figure 12. Putatively polysaccharide biosynthesis pathways in Seep-SRB1a** | **13** |
| **Supplementary Figure 13. Putatively polysaccharide biosynthesis pathways in Seep-SRB1g** | **14** |
| **Supplementary Figure 14. Putatively polysaccharide biosynthesis pathways in Seep-SRB2** | **15** |
| **Supplementary Figure 15. Putatively polysaccharide biosynthesis pathways in HotSeep-1** | **16** |
| **Supplementary Figure 16. Presence of extracellular contractile injection systems (eCIS) in ANME and SRB.** | **17** |
| **Supplementary Figure 17. Phylogeny of afp10, the spike protein from the extracellular contractile injection system.** | **18** |
| **Supplementary Figure 18. Phylogeny of afp11, a base plate protein from the extracellular contractile injection system.** | **19** |

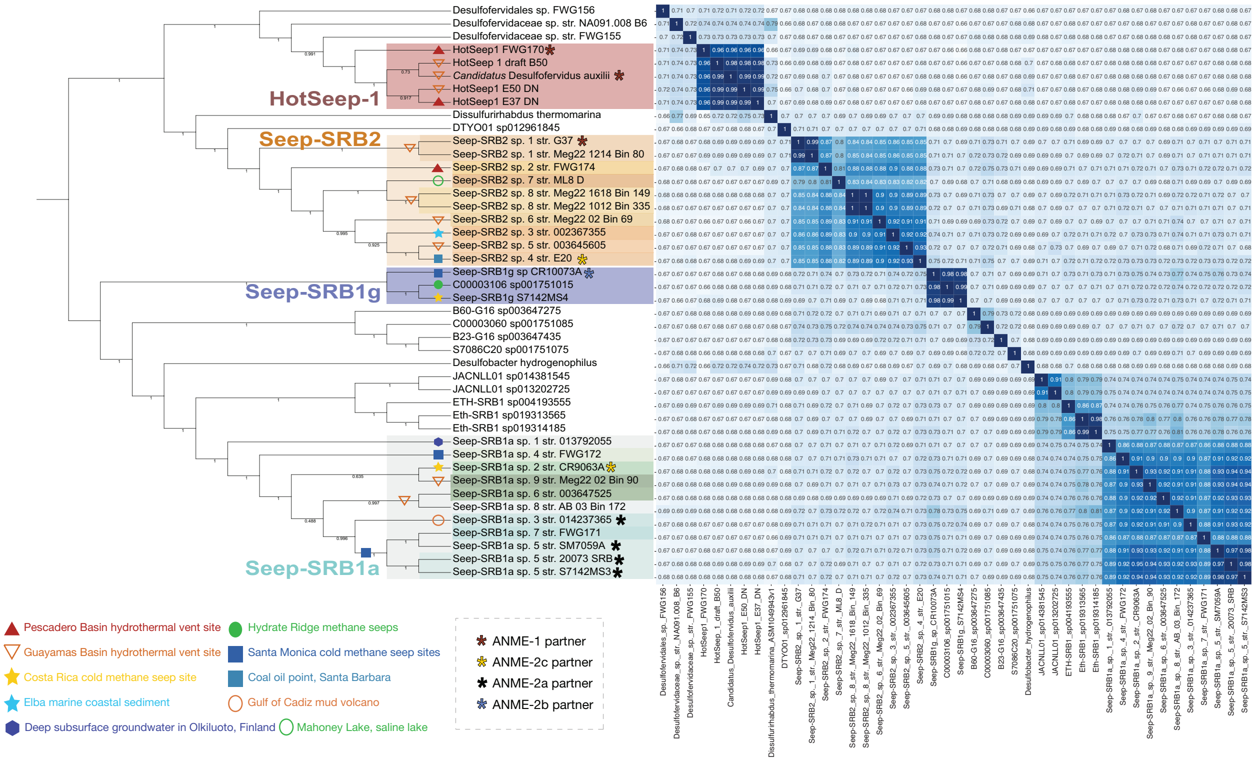


**Supplementary Figure 1. Taxonomic separation of syntrophic sulfate reducing bacteria using average nucleotide identity**. The average nucleotide identity of genomes from each clade of the syntrophic sulfate reducing bacteria and some related bacteria were computed using the PyANI program available through Anvi'o. The different clades of syntrophic SRB – HotSeep1, Seep-SRB2, Seep-SRB1a and Seep-SRB1g are colored according to the attached legend. The Seep-SRB1a genomes in particular are differently colored depending on whether they partner

ANME-2a or ANME-2c respectively. The geographic location from which each genome was extracted is indicated on each node or clade in the tree.

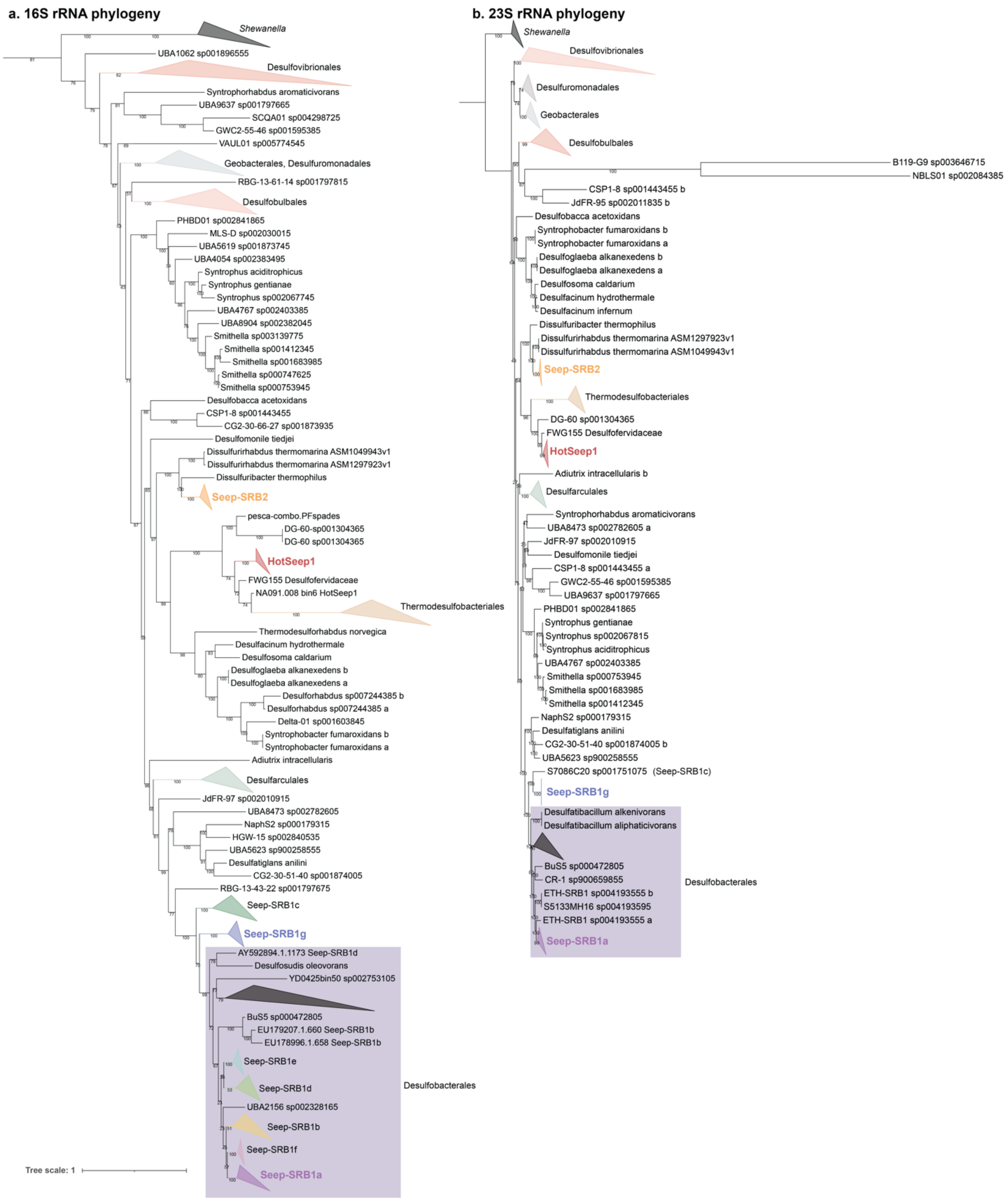


**Supplementary Figure 2. Comparison of 16S rRNA and 23S rRNA phylogeny of organisms from the phylum Desulfobacterota.** 16S rRNA and 23S rRNA sequences were extracted from all organisms from the phylum Desulfobacterota available in GTDB release 95, and from syntrophic SRB. These sequences were aligned using MUSCLE and a tree was inferred using IQTREE2. Hot-Seep1 is placed adjacent to the order Thermodesulfbacteriales in both these trees.

**a. RpoB phylogeny**

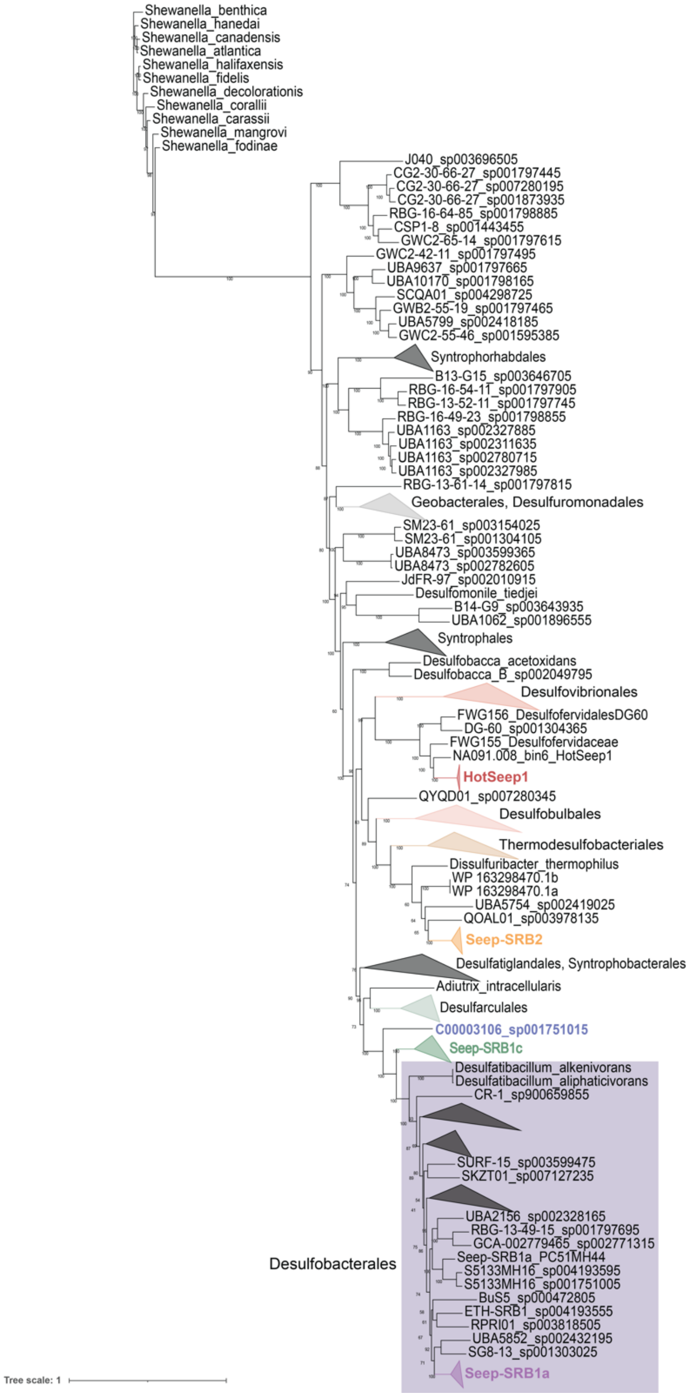


**Supplementary Figure 3. Phylogeny of RpoB from organisms within the phylum Desulfobacterota.** Sequences of RNA Polymerase, subunit B were extracted from all organisms from the phylum Desulfobacterota available in GTDB release 95, and from syntrophic SRB using BLASTP with an e-value cut-off of e-30 and appropriate query sequences. The sequences were confirmed to RpoB by manual inspection of a multiple sequence alignment generated using

MUSCLE and the tree was inferred using IQTREE2. In this tree, Hot-Seep1 is found adjacent to the order Desulfovibrionales.

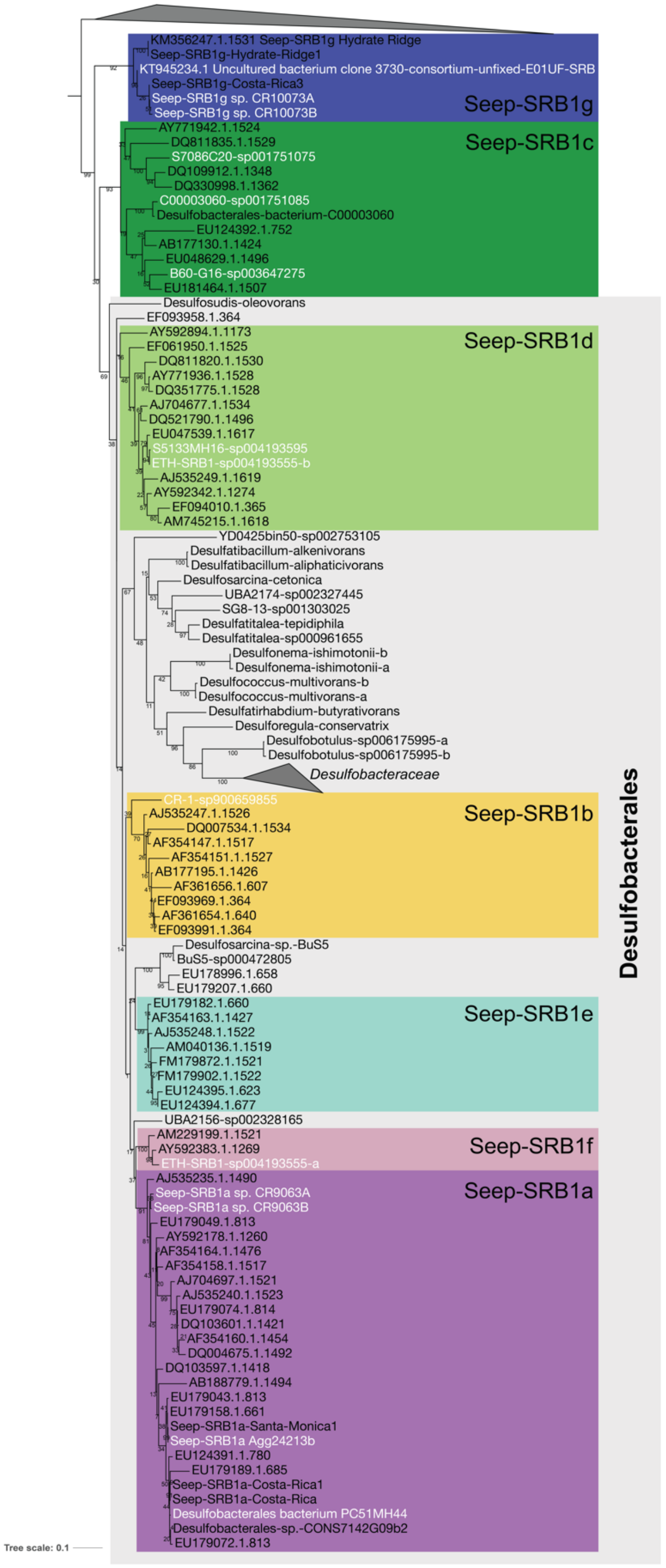

**Supplementary Figure 4. Placement of various Seep-SRB1 clades within the phylum Desulfobacterota.** A phylogenetic tree of full length 16S sequences from various Seep-SRB1 clades including the original 16S rRNA sequences used to define the Seep-SRB1(a-f) clades.

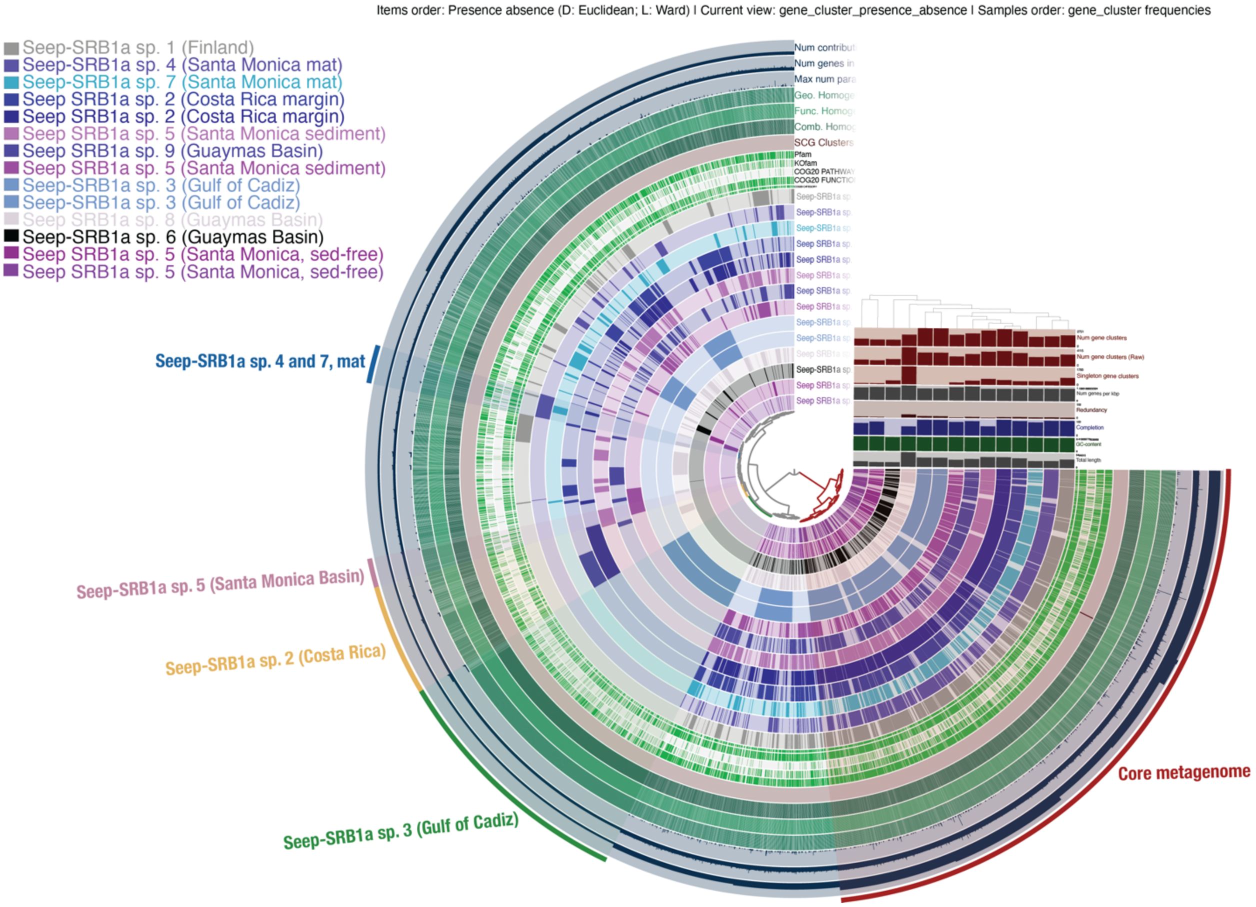


**Supplementary Figure 5. Pan-genome analysis of Seep-SRB1a metagenomes.** Fourteen genomes from nine Seep-SRB1a species were analyzed using the Anvi’o pan-genome analysis pipeline. Five gene cluster bins were annotated based on genes that were identified as part of the core metagenome, unique to Seep-SRB1a sp. 2, Seep-SRB1a sp. 3, Seep-SRB1a sp. 5 and from the Seep-SRB1a sp. 4 and 7.

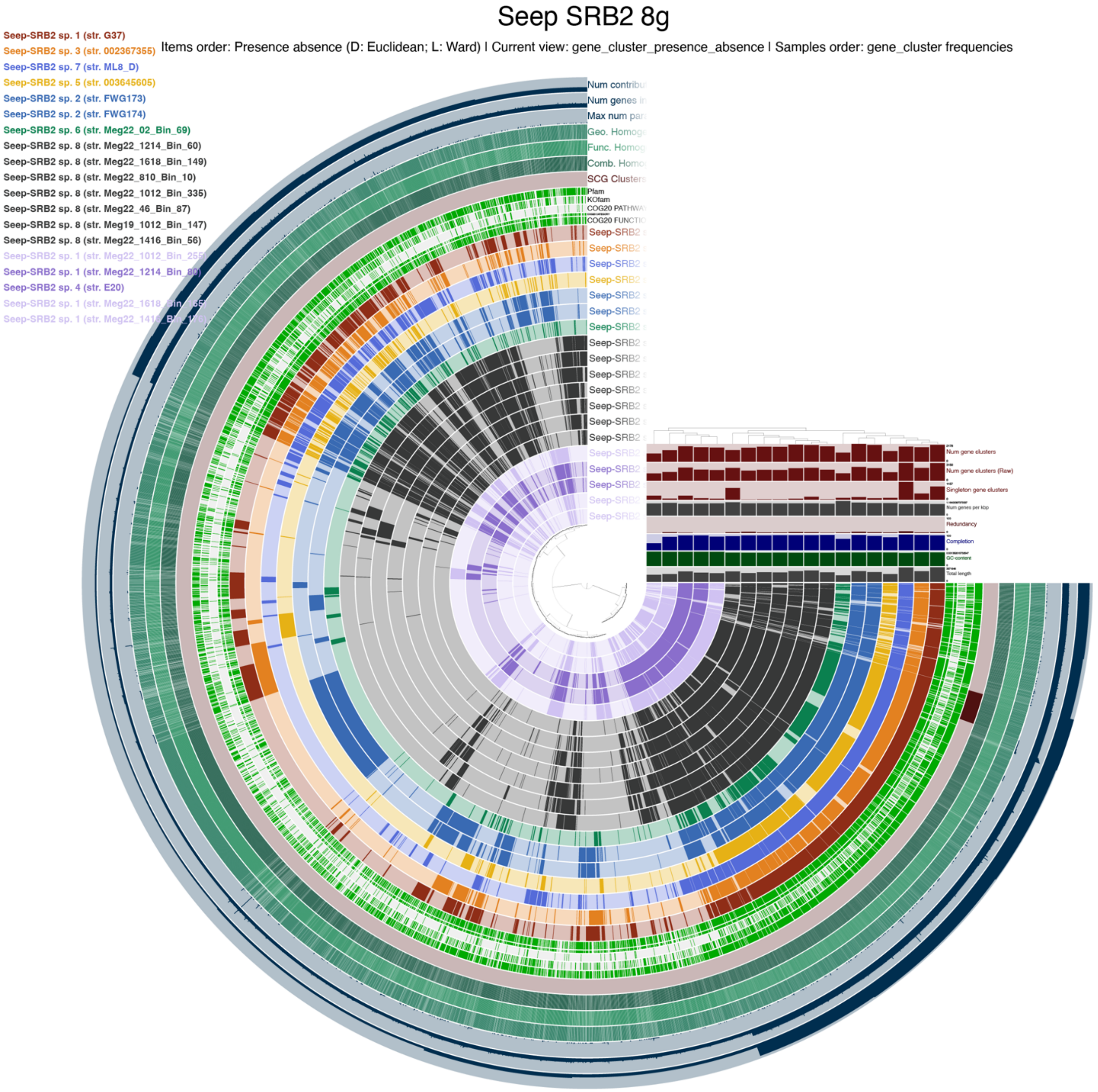


**Supplementary Figure 6. Pan-genome analysis of Seep-SRB2 metagenomes.** Fourteen genomes from nine Seep-SRB1a species were analyzed using the Anvi'o pan-genome analysis pipeline. Three gene cluster bins were annotated based on genes that were identified as part of the core metagenome, present in Seep-SRB2 sp. 1 and absent in Seep-SRB2 sp.1

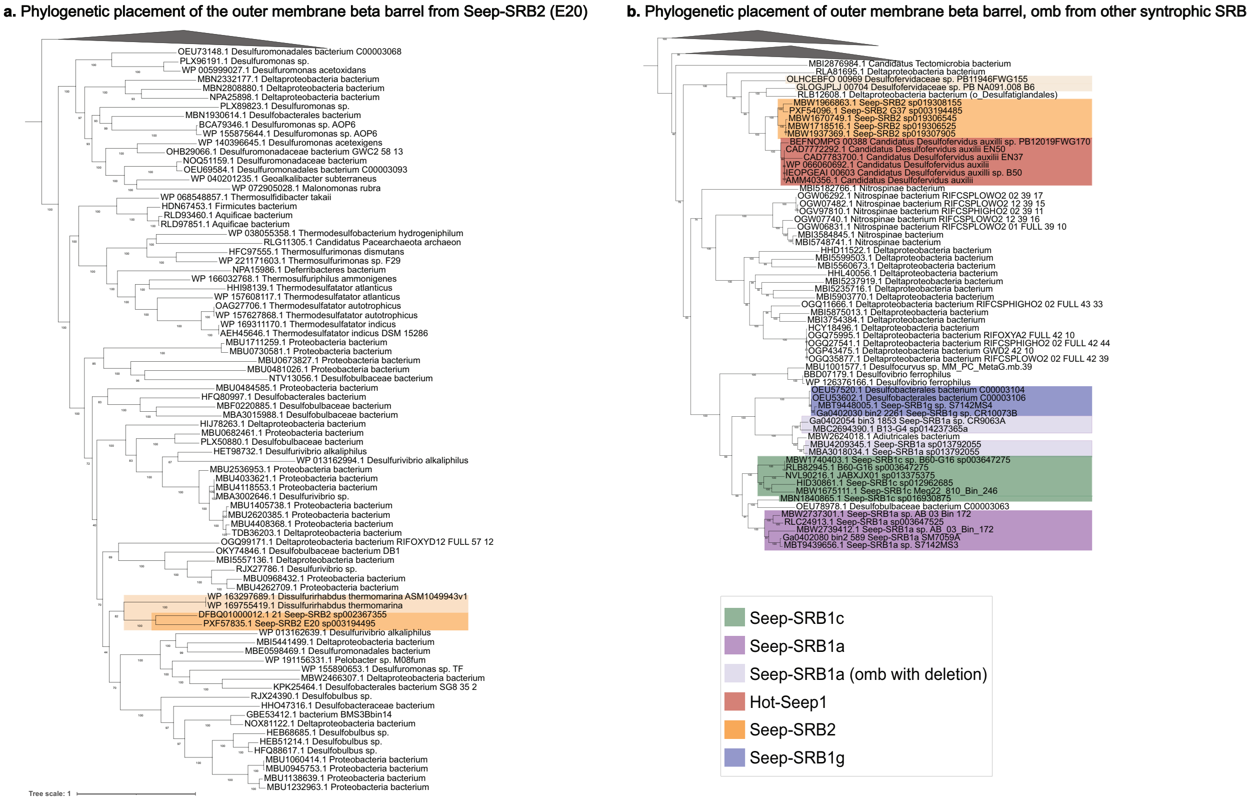


**Supplementary Figure 7. Phylogenetic placement of the outer membrane beta barrel, OetI from the putative DIET cluster**. A multiple sequence alignment, **Supplementary multiple sequence alignment MSA2** of the OetI protein sequences extracted from the genomes of syntrophic SRB and the NCBI database was generated using MUSCLE. This alignment was used to infer a phylogenetic tree using IQ-Tree2 and visualized on the iTOL web server. a. The phylogenetic placement of OetI from E20 Seep-SRB2 next to OetI from Thermodesulfobacteria and *Dissulfurirhabdus thermomarina* demonstrates that it was possibly vertically acquired from a gene transfer that was ancestral to the Seep-SRB2 and then vertically transferred. b. The phylogenetic placement of Seep-SRB1a and Seep-SRB1g OetI suggests that they are related. Additionally, the placement of OetI from G37 Seep-SRB2 next to OetI from Desulfofervidales, suggests that the Seep-SRB2 partner of ANME-1 acquired its DIET cluster from HotSeep1.

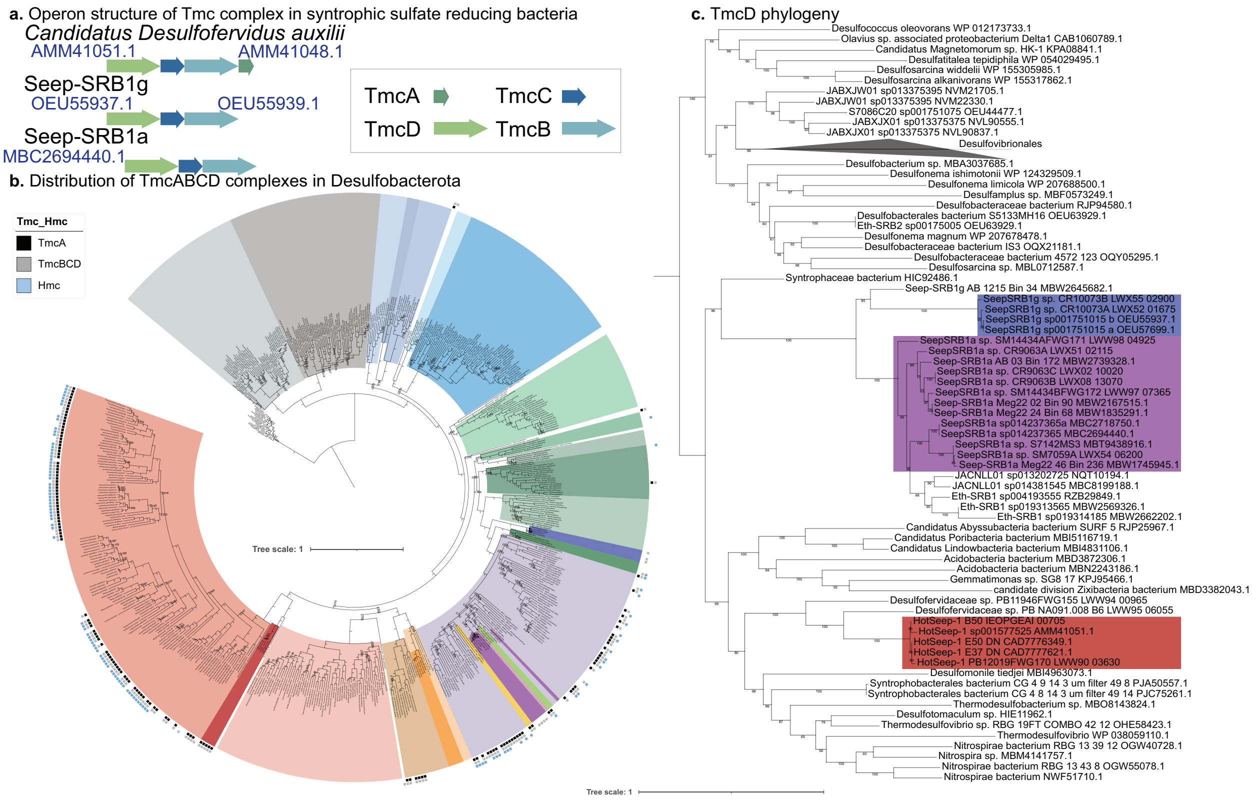


**Supplementary Figure 8. The Tmc complex in Seep-SRB1a and Seep-SRB1g are divergent**. A. The operons containing Tmc in *Candidatus Desulfofervidus auxilii* (Hot-Seep1), Seep-SRB1g and Seep-SRB1a show that TmcA is present in the former operon while it is missing in the latter two. B. The distribution of Tmc is mapped on to the phylum Desulfobacterota, showing that it is common in the order Desulfobacterales and Desulfovibrionales. C. Phylogeny of TmcD demonstrates that the Tmc complex in Seep-SRB1a and Seep-SRB1g cluster together and appear to be different from the Tmc complex in other organisms from the orders Desulfobacterales and C00003060.

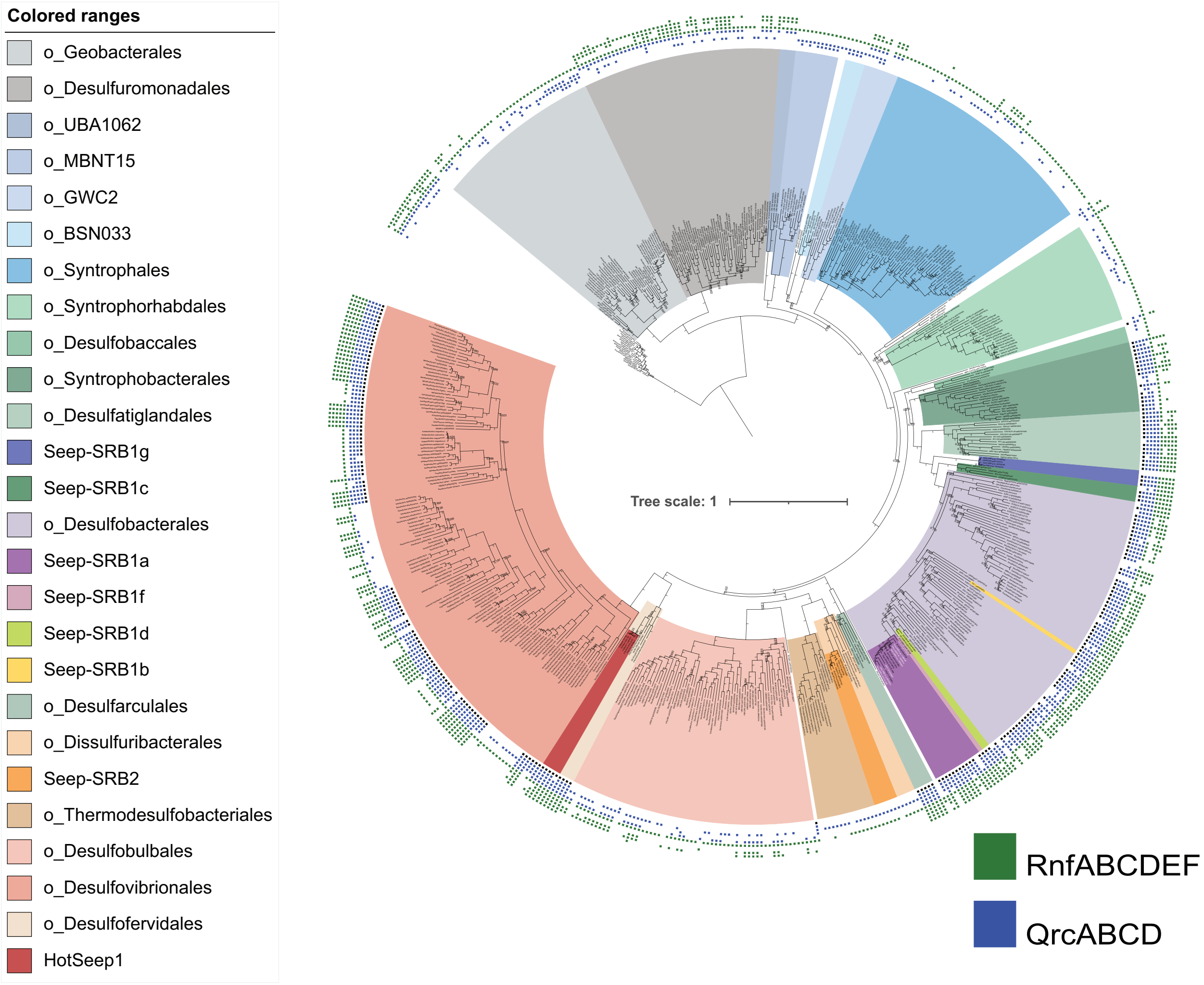


**Supplementary Figure 9. Distribution of QrcABCD and RnfABCDEG in Desulfobacterota.** The presence and absence of Qrc and Rnf was demonstrated across the Desulfobacterota using BLASTP searches of different query sequences of these complexes. Both these complexes are absent from the orders Desulfobulbales, Thermodesulfobacteriales and Dissulfuribacteriales.

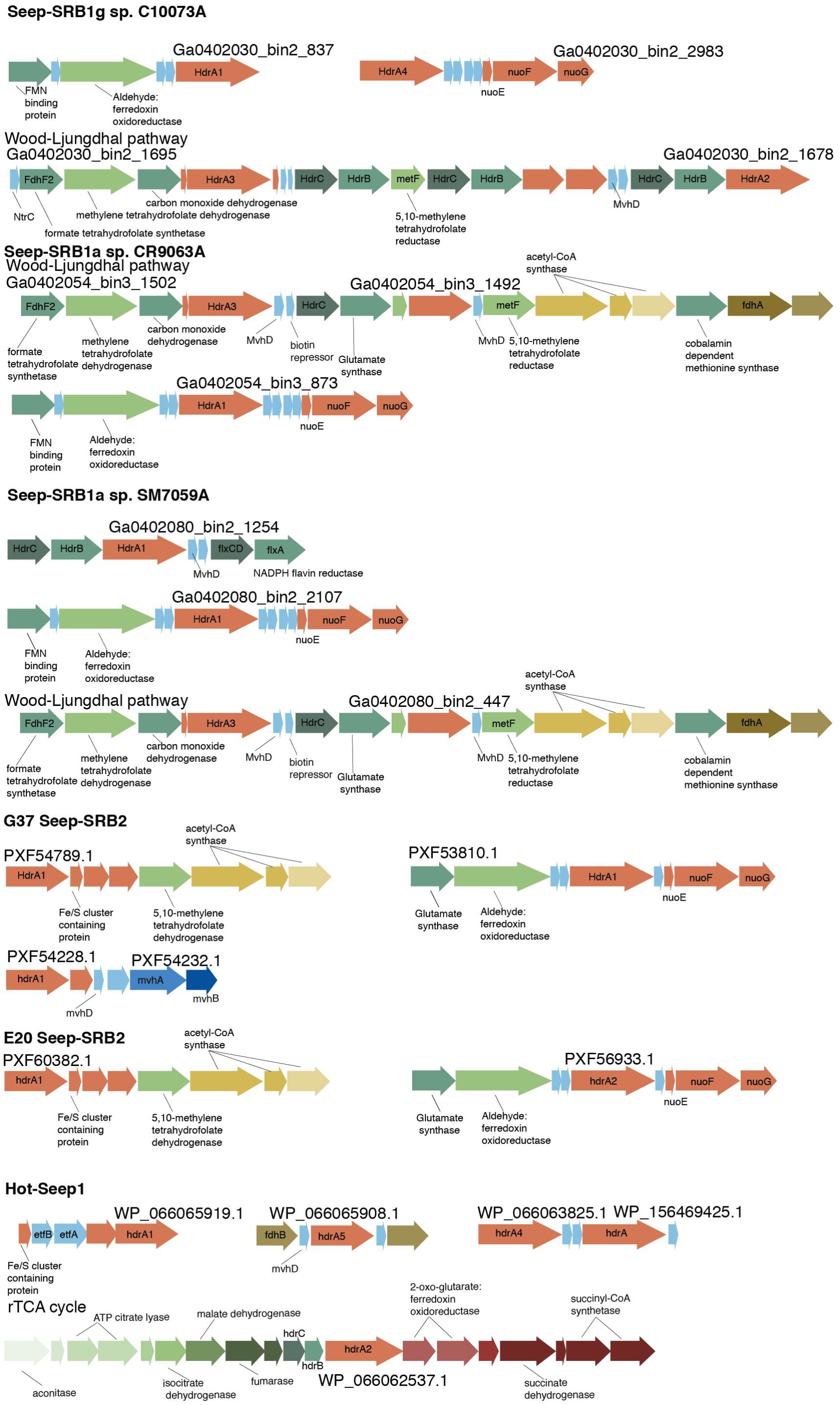


**Supplementary Figure 10. Gene neighborhoods of various HdrA containing complexes and carbon fixation pathways in syntrophic SRB.** HdrA homologs were identified in the four syntrophic SRB clades with the following gene neighborhoods. Multiple HdrA homologs were identified adjacent to the carbon fixation pathways in syntrophic SRB, specifically Seep-SRB1a and Seep-SRB1g.

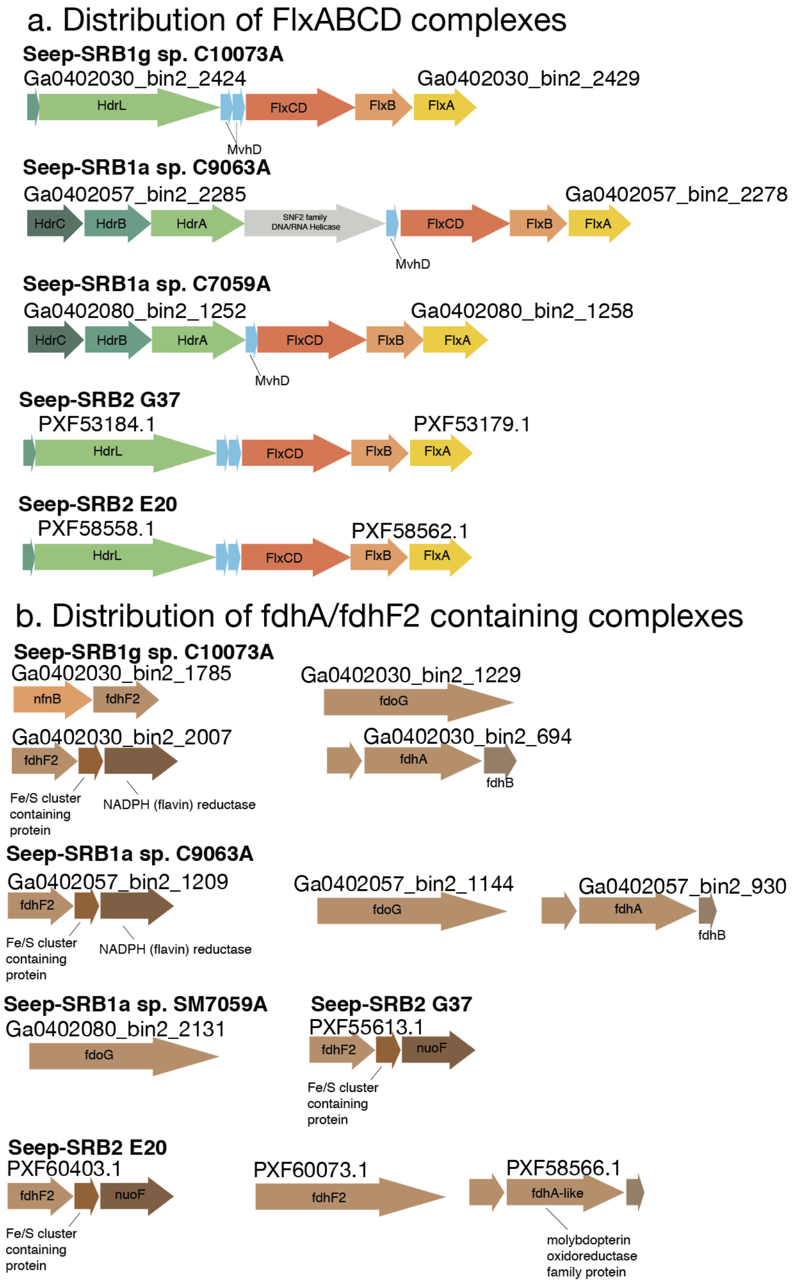


**Supplementary Figure 11. Operons of Flx-Hdr complexes and gene neighborhood of putative formate utilizing proteins.** a. Flx-Hdr complexes which recycle electrons between NADH, ferredoxins and DsrC are found in Seep-SRB1g, Seep-SRB1a and Seep-SRB2. b. Many putative formate utilizing proteins are found in Seep-SRB1g, Seep-SRB1a and Seep-SRB2. In Seep-SRB1g and Seep-SRB1a, periplasmic formate dehydrogenases (fdhAB) are found. fdhA domains as identified here are typically found in the periplasm and have a respiratory function while fdhF2 are typically cytoplasmic.

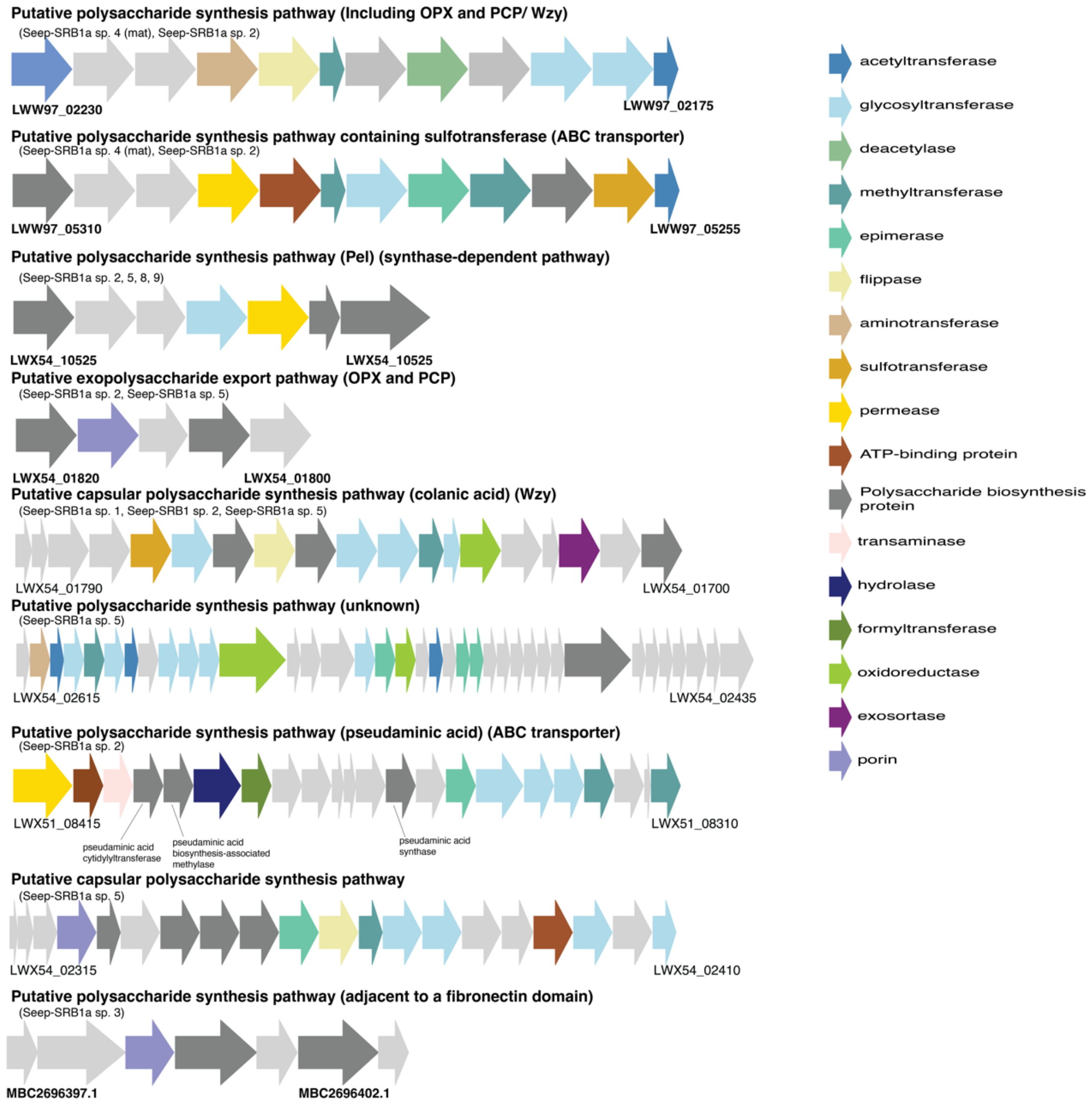


**Supplementary Figure 12. Putative polysaccharide biosynthesis pathways in Seep-SRB1a.**

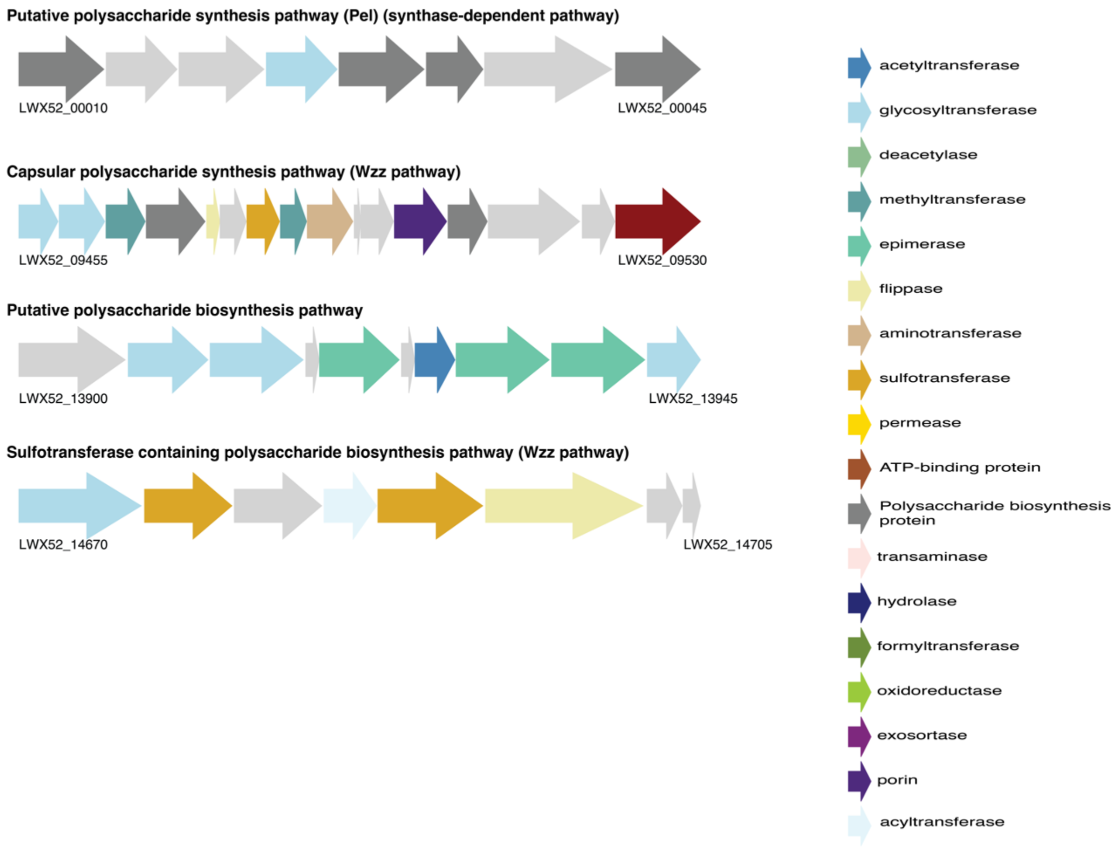


**Supplementary Figure 13. Putative polysaccharide biosynthesis pathways in Seep-SRB1g.**

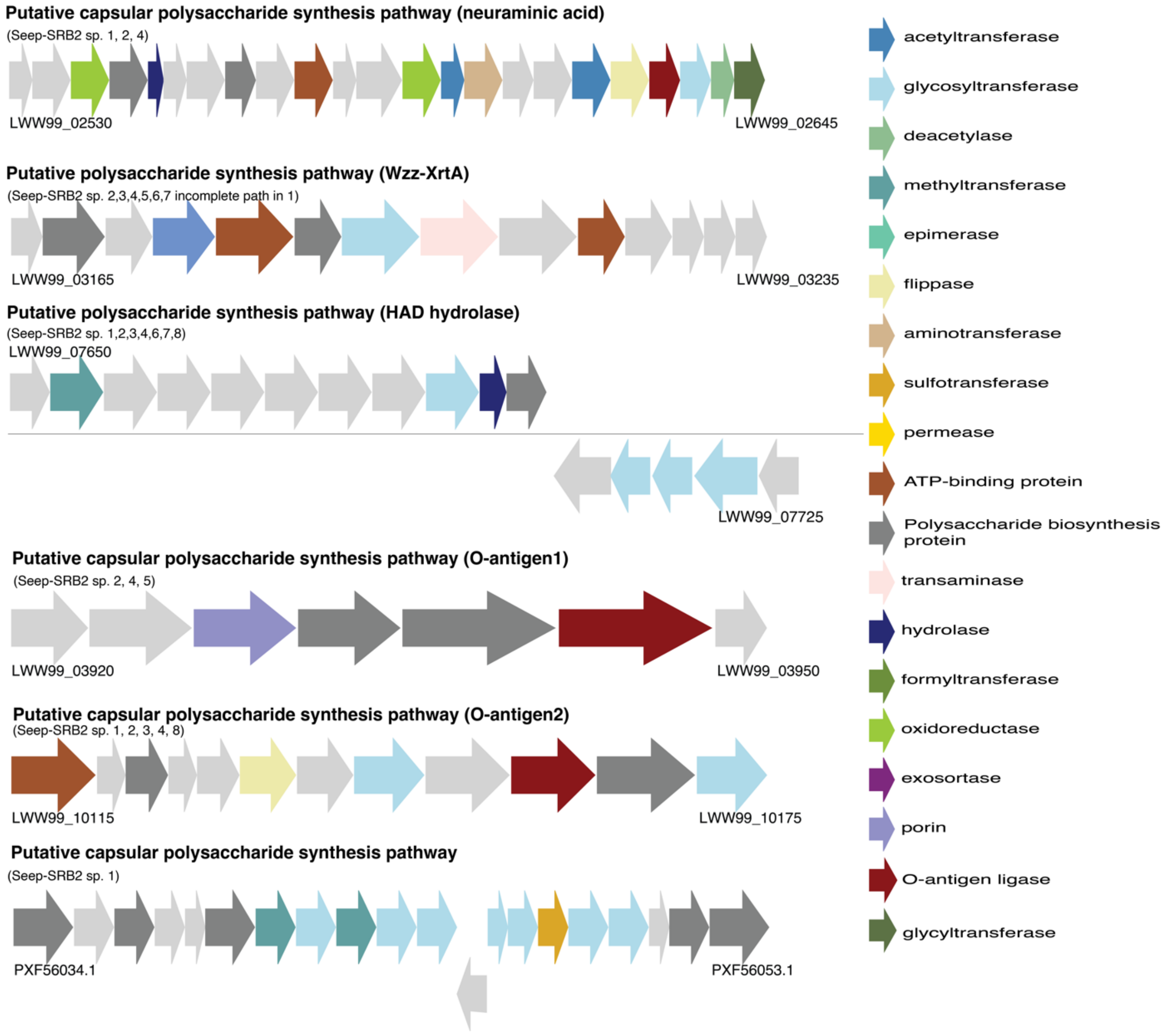


**Supplementary Figure 14. Putative Polysaccharide biosynthesis pathways in Seep-SRB2.**

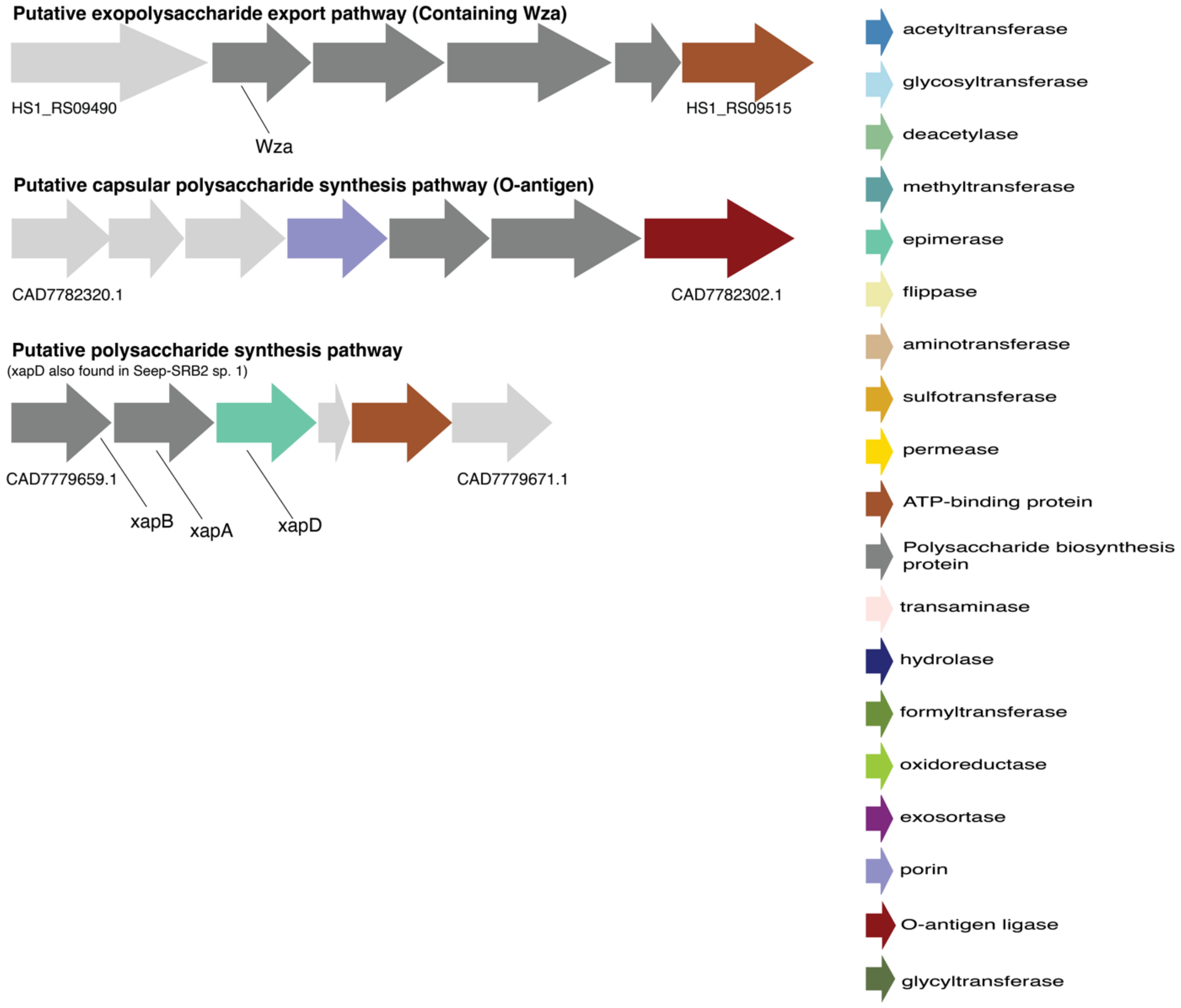


**Supplementary Figure 15. Putative polysaccharide biosynthesis pathways in HotSeep-1.**

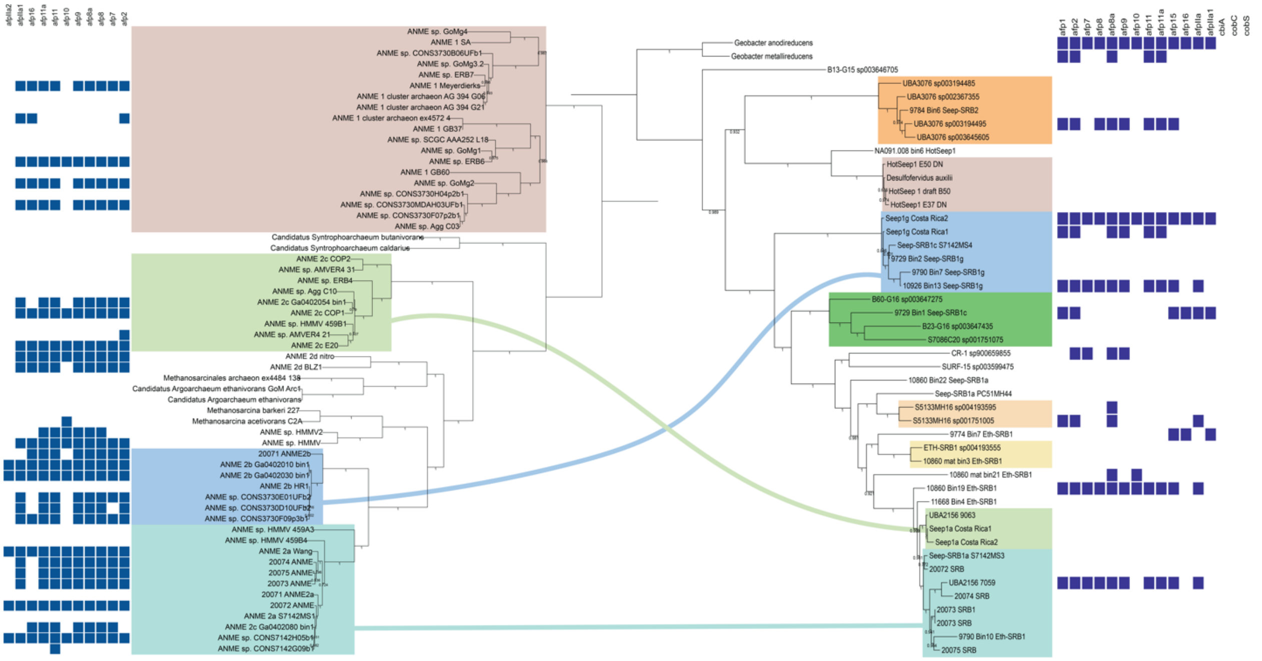


**Supplementary Figure 16. Presence of extracellular contractile injection systems (eCIS) in ANME and SRB.** The presence of eCIS conduits in ANME and SRB was identified using BLASTP and dbeCIS[cite Chen et al. Cell Reports]. While the eCIS clusters are widely distributed in ANME-2a and ANME-2b, they are only sparsely distributed in ANME-1. They are only present in the Seep-SRB1g species found in Costa Rica and one of the Seep-SRB1a species from Santa Monica Basin.

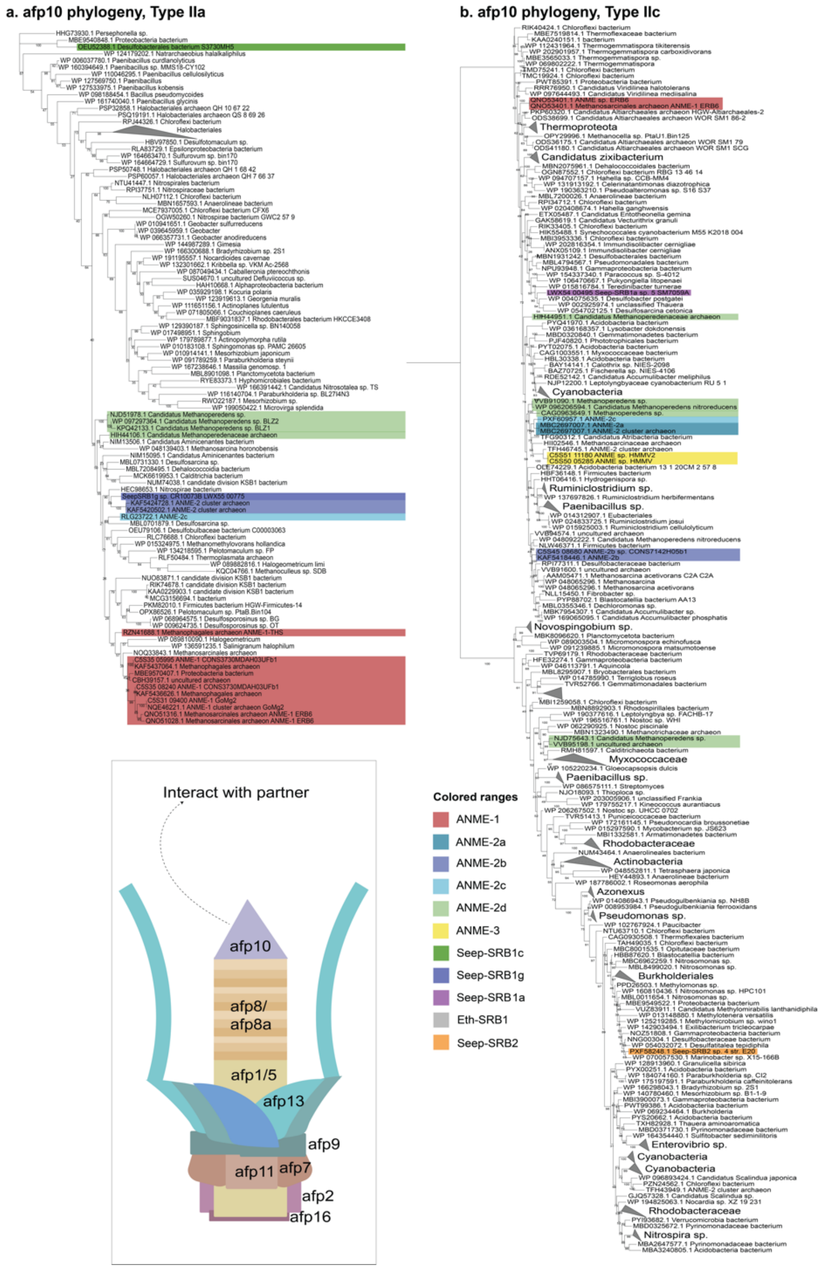


**Supplementary Figure 17. Phylogeny of afp10, the spike protein from the extracellular contractile injection system.** Afp10 is the PAAR-domain containing protein that typically interacts with the target organism of the eCIS. Afp10 sequences were extracted from ANME and SRB using BLASTP and dbeCIS. These sequences were then used as queries to repeatedly search and identify the closest homologs from the NCBI database. A sequence alignment was then made using MUSCLE, manually inspected and filtered, and the tree was inferred using RAxML. Seep-SRB1a sequences are related to other eCIS sequences from Desulfobacterales while Seep-SRB2 afp10 and Seep-SRB1g afp10 sequences do not cluster with evolutionarily related bacteria.

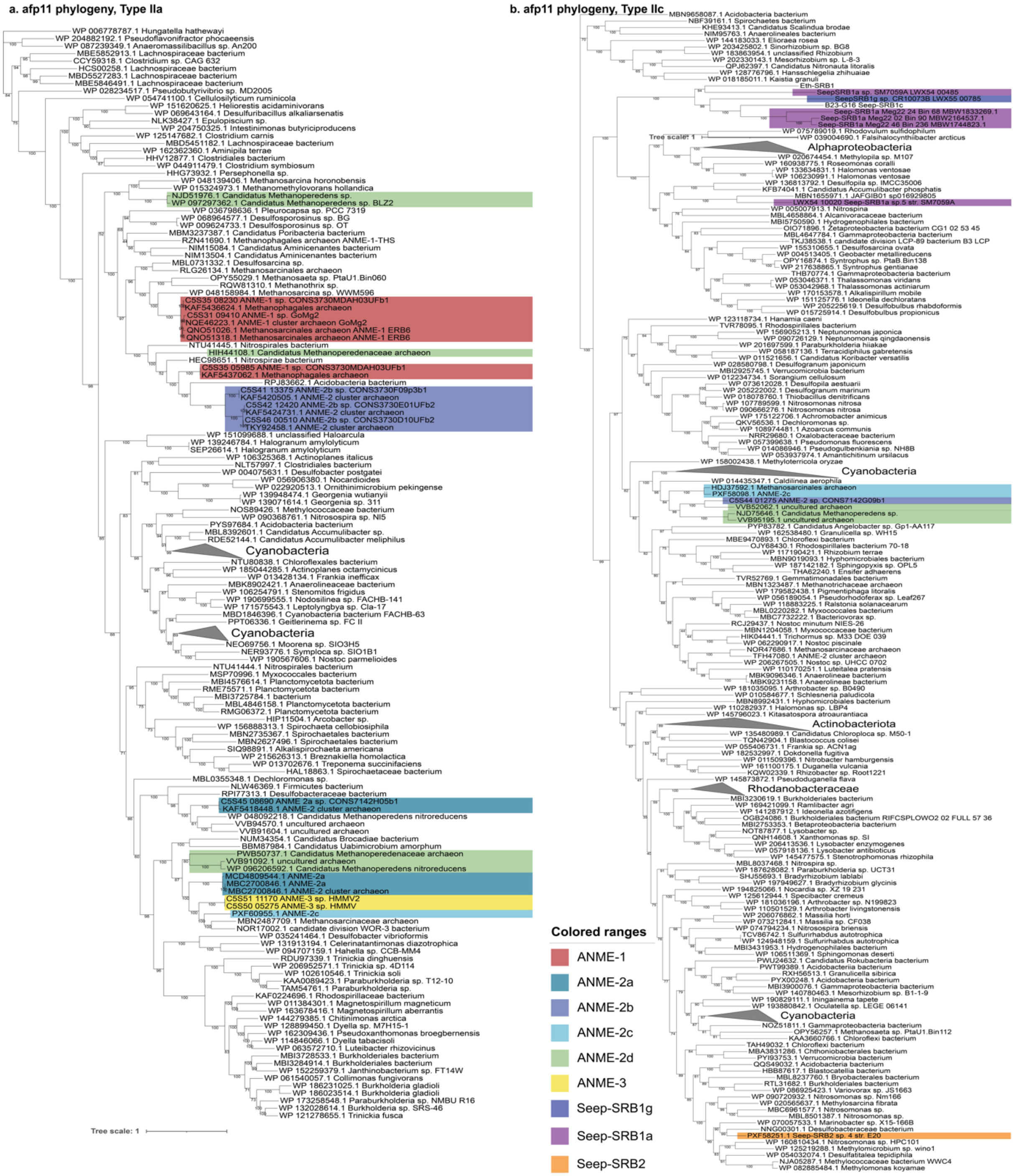


**Supplementary Figure 18. Phylogeny of afp11, a base plate protein from the extracellular contractile injection system.** Afp11 belongs to the baseplate of the eCIS and does not interact directly with the target organism. Afp11 sequences were extracted from ANME and SRB using BLASTP and dbeCIS. These sequences were then used as queries to repeatedly search and identify the closest homologs from the NCBI database. A sequence alignment was then made using MUSCLE, manually inspected and filtered, and the tree was inferred using RAxML. Seep-SRB1a sequences are related to other eCIS sequences from Desulfobacterales while Seep-SRB2 afp10 and Seep-SRB1g afp10 sequences do not cluster with evolutionarily related bacteria.
